# Emergence and stabilisation of a neo-Y chromosome in nematode species with rare males

**DOI:** 10.1101/2025.07.25.666552

**Authors:** Brice Letcher, Lewis Stevens, Nathanaëlle Saclier, Andrew Hsiao, Manuela Kieninger, Eva Wenger, Caroline Launay, Mark Blaxter, Marie Delattre

**Affiliations:** LBMC, ENS de Lyon and CNRS, Lyon, France; Tree of Life, Wellcome Sanger Institute, Cambridge, United Kingdom; ISEM, Université Montpellier, France

## Abstract

How do sex chromosomes evolve in the transition to asexuality? So far, species that depart from canonical sexual reproduction - for example, parthenogens with rare sex, or species where the paternal genome is set aside - have been found to carry either no sex chromosomes, or sex chromosomes but no male-specific sex chromosome (i. e., no Y). Here we reveal that, in *Mesorhabditis* nematodes, a new Y chromosome emerged once, from sexual ancestors, in species that have transitioned into an unconventional mode of reproduction called autopseudogamy. In this reproductive system, females produce clonal females plus rare (∼10%) males that are needed for fertilisation, but that do not contribute to the female genome. Analysing the Y chromosomes of two autopseudogamous species, we found high levels of degeneration, most likely due to loss of recombination, and two additional conserved features: i) they accumulated male-beneficial genes, and ii) they display a strong fertilisation drive, in that mainly Y-bearing sperm fertilise female oocytes. Both features are likely evolutionarily favourable in the context of autopseudogamy. Our results suggest that male-specific chromosomes can still be maintained in systems with rare, and possibly ‘genetically useless’, males.

## Introduction

Sex chromosomes evolve in many fascinating ways because they are subjected to unique evolutionary forces. They often carry sex determining loci and/or rapidly-evolving sexually antagonistic genes, whose linkage can select for loss of recombination in the heterogametic sex (XY in XX/XY systems, and ZW in ZZ/ZW systems), in turn leading to differentiation and degeneration of the Y or Z (1–3). Y chromosomes are often the targets of selfish drivers reducing their transmission rate during male meiosis, creating an arms race on the sex chromosomes to restore equal sex ratios (4). In some taxa, sex chromosomes have been relatively stable over millions of years (e.g. therian mammals (5)), while in others, sex chromosomes have been prone to rapid turnovers, fusions with other chromosomes, changes in the heterogametic sex and transitions to an XX/XO system following the loss of a Y chromosome (e.g. fish, amphibians, reptiles, dipteran insects (6,7)).

Most knowledge on the evolution of sex chromosomes comes from the study of sexual species displaying nearly equal sex ratio and little is known about the fate and evolution of sex chromosomes as reproductive modes depart from canonical sexuality. Currently, known species that have lost sex entirely - called parthenogens - have no recognisable sex chromosomes (e.g. bdelloid rotifers (8,9), dandelions (10)). Other intermediate cases of partial asexuality have been studied, including parthenogens producing rare males, cyclical parthenogens and species producing males that transmit only their mother’s genome after paternal genome elimination. In all these cases, sex is either determined via ploidy (e.g. insects undergoing paternal genome elimination (11,12)), via environmental determinants (e.g. *Daphnia* crustaceans (13)), via XX/XO sex chromosomes (e.g. *Timema* stick insects (14), pea aphids (15)), or ZZ/ZW sex chromosomes, whereby females are the heterogametic sex (e.g. *Artemia* shrimp (16)). In other words, no male-specific Y chromosome has been found when males are rarely produced or do not transmit their genome. These examples suggest that Y chromosomes may easily be lost when the effective population size of males becomes too small, or that pseudosexuality emerges preferentially in clades with no Y chromosome.

Here, we study sex chromosome evolution in the nematode genus *Mesorhabditis,* in which a peculiar mode of reproduction called autopseudogamy has evolved once, from sexual ancestors (17). Autopseudogamous species feature only 2-14% males in the population that are essential for reproduction, as male sperm is needed to activate the eggs. In ∼90% cases however, the sperm DNA is set aside after fertilisation and eggs develop into females from the maternal genome only (i.e. gynogenesis). The remaining embryos are produced after regular mixing of the parental genomes, i.e. amphimixis, and in this case, give rise to sons only. We previously found that one autopseudogamous species, *M. belari*, is XX/XY, and that only sons are produced following amphimixis due to a strong fertilisation drive in favour of the Y-bearing sperm (18) (Fig. 1A).

**Fig. 1.**
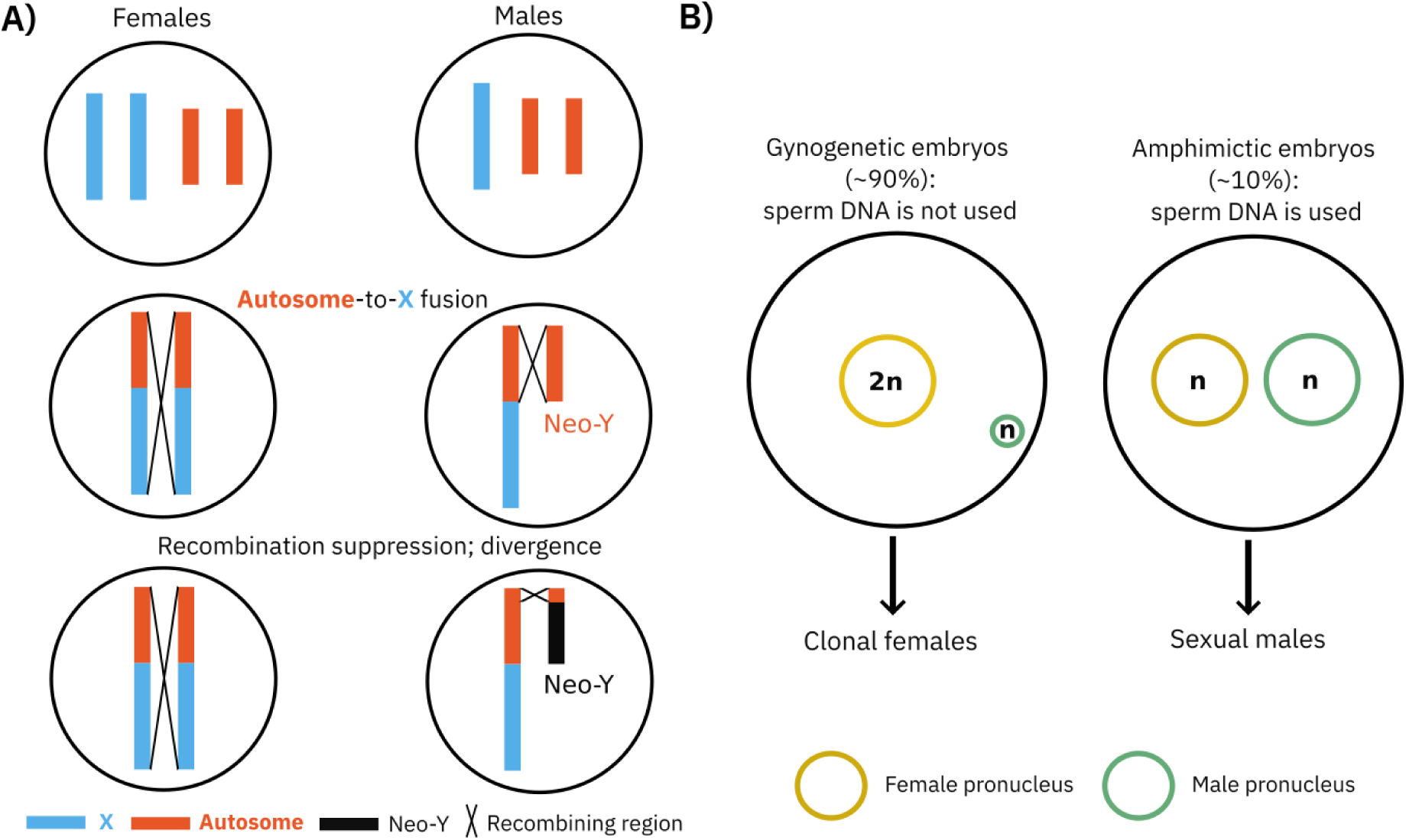
Introductory concepts: autopseudogamy and X-autosome fusions. **A) Autopseudogamy**. In this mode of reproduction, found in several species of *Mesorhabditis*, females produce two types of embryos during their lifetime: gynogenetic embryos (90% of embryos; left-hand cell), in which oocytes do not use the sperm DNA (green pronucleus) and undergo a modified meiosis to maintain diploidy (2n inside the yellow female pronucleus). These give rise to asexual clonal females. The remaining embryos are amphimictic (10% of embryos; right-hand cell), in which oocytes undergo regular meiosis and mix with the haploid sperm DNA. In this case, only males are produced. We previously found that in one species, *M. belari*, this is due to a strong fertilisation drive in favour of Y-bearing sperm (18). **B) Neo-Y emergence by X-autosome fusion**. A female diploid for the X chromosome (blue) and one pair of autosomes (orange) is shown on the left, and a male diploid for the autosome pair and haploid for the X is shown on the right. In this XX/X0 system, if an autosome fuses to the X, individuals diploid for the fusion become females, while individuals haploid for the fusion become males. The unfused autosome thus forms a neo-Y chromosome *de facto*. In males, the unfused autosome may persist and keep recombining with its fused counterpart. If recombination gradually ceases however, the unfused autosome diverges specifically in males and becomes a ‘true’ Y chromosome (bottom-right cell).

Here, we tested whether a Y chromosome is specifically associated with autopseudogamy. In nematodes, most species are XX/X0 - including *C. elegans,* part of the same family as *Mesorhabditis* (Rhabditidae). Neo-Y chromosomes can easily arise in XX/X0 systems through X-autosome fusions (Fig. 1B) (19), but neo-Y chromosomes have so far been rarely identified in nematodes, in a few sexual species only (20–22). We thus explored the evolutionary origin of the Y chromosome in the autopseudogamous (hereafter, for simplicity, referred to as pseudosexual) species of *Mesorhabditis*, and the possible reasons for its existence and maintenance.

## Results

### Sexual species are XX/X0 and pseudosexual species are XX/XY in *Mesorhabditis*

We have previously established that pseudosexuality arose a single time in *Mesorhabditis* (17) and that one pseudosexual species, *M. belari*, has male-specific genome regions clearly consistent with a Y chromosome (18). To establish the origin of the Y chromosome, we assembled the genome to high contiguity of four species spanning phylogenetic diversity of *Mesorhabditis*, using long-read DNA-Seq data plus Hi-C for scaffolding: two sexual species, *M. spiculigera*, and *M. longespiculosa*, the pseudosexual species *M. monhystera,* and an improved assembly for the pseudosexual species *M. belari* (Fig. 2A).

**Fig. 2.**
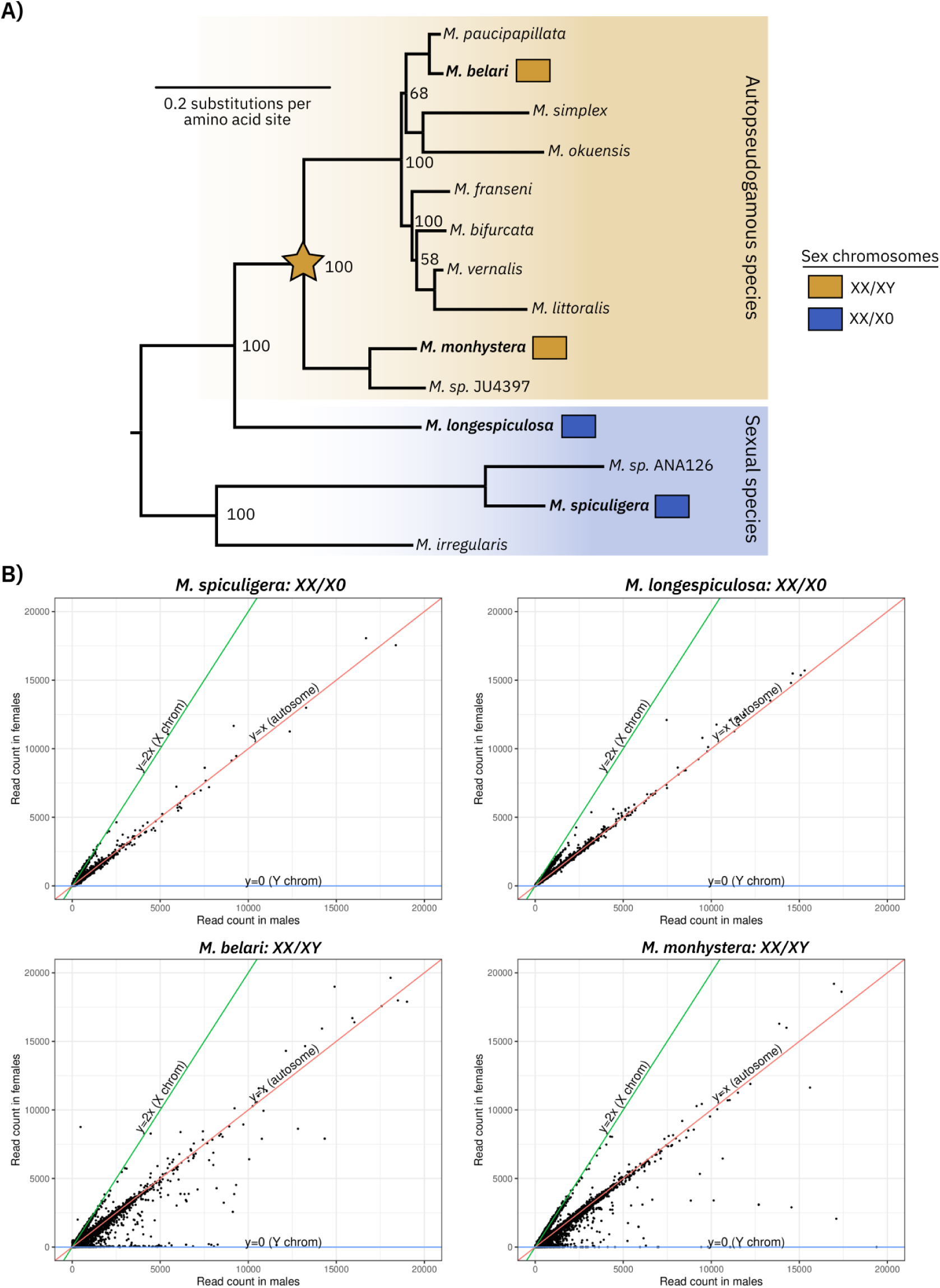
Co-occurrence of a neo-Y chromosome and pseudosexual reproduction in *Mesorhabditis*. **A)** Phylogeny of *Mesorhabditis* species, built from conserved protein-coding genes found across all species (see Methods). Numbers at the nodes indicate the percentage of trees supporting each node, out of 1000 bootstrapped trees. Coloured boxes show modes of reproduction (as identified in the lab (17)), with autopseudogamous species shaded in orange and sexual species in blue. These clearly support a single emergence of autopseudogamy (that we refer to as pseudosexuality), marked with an orange star on the phylogeny. The four species in bold are those for which we identified the sex chromosomes using assembled genomes and male/female DNA-Seq: the data are consistent with a single emergence of a neo-Y chromosome on the branch indicated by the orange star. **B)** Identification of sex chromosomes using male/female DNA-Seq. Sequencing reads were aligned to predicted genes in the genomes of the four focal species (*M. belari*, *M. monhystera*, *M. longespiculosa*, *M. spiculigera*): genes with the same count in males and females are autosomal (red curve), twice the coverage in females are on the X (green curve), and coverage in males but no coverage in females are on the Y (blue curve). Genes consistent with a Y chromosome are only seen in the two pseudosexual species (*M. belari* and *M. monhystera*).

We obtained N50 values of 514kbp-1.8Mbp, for genome sizes 140-240Mbp, and BUSCO completeness values of 81-84%. We note that our assemblies are not chromosome-level, however, in large part due to unbridged long repeat arrays (see Methods for details).

To test for the presence of sex chromosomes, we generated short-read DNA-Seq data from separated pools of males and females that we mapped to predicted genes in the genomes. We focussed on genes as they are easier to map to, due to lower repeat content (see Methods for details). After normalising for sequencing depth, we compared male and female coverages: genes with twice the coverage in females should be on the X, and coverage only in males should be on the Y. Genes with equal coverage in both should be on autosomes (or on a portion of the Y that has not yet diverged from the X - we return to this later). This approach clearly demonstrated that male-specific genes were identified in the two pseudosexual species only, whereas all species had X-like genes (Fig. 2B). From these results we concluded that the two sexual *Mesorhabditis* species are XX/X0 while the two pseudosexual species are XX/XY. Thus, at the phylogenetic level, sexual *Mesorhabditis* species have maintained the ancestral XX/XO sex determining system of other species in the *Rhabditidae* family (23,24) whereas a Y chromosome co-occurs with pseudosexuality (Fig. 2A, yellow star).

### Evolution of the X chromosome in *Mesorhabditis*: turnover and fusions

To understand how the sex chromosomes of *Mesorhabditis* evolved, we assigned entire assembled scaffolds to X, Y or autosomal status, by comparing male/female coverage ratios along entire scaffolds. As genes are easier to map to, we also used male/female coverage ratios along genes to check for consistency. As an example, a X-assigned scaffold for *M. monhystera* is shown in Fig. 3: almost all genes are clearly X-assigned (Fig. 3A), and the entire scaffold has diploid coverage in females and haploid coverage in males, consistent with an X chromosome (Fig. 3B). Almost all large assembled scaffolds (>100kbp) could be confidently assigned in this way, and when not, visual inspection was used to resolve the scaffold’s status (see Methods). For the two pseudosexuals, we obtained two X-assigned scaffolds, while for the sexuals *M. spiculigera* and *M. longespiculosa*, we obtained 6 and 20, respectively. The latter are more fragmented overall, likely due to higher heterozygosity in sexual species, and due to lack of Hi-C data for assembly specifically for *M. longespiculosa* (see Methods).

**Fig. 3.**
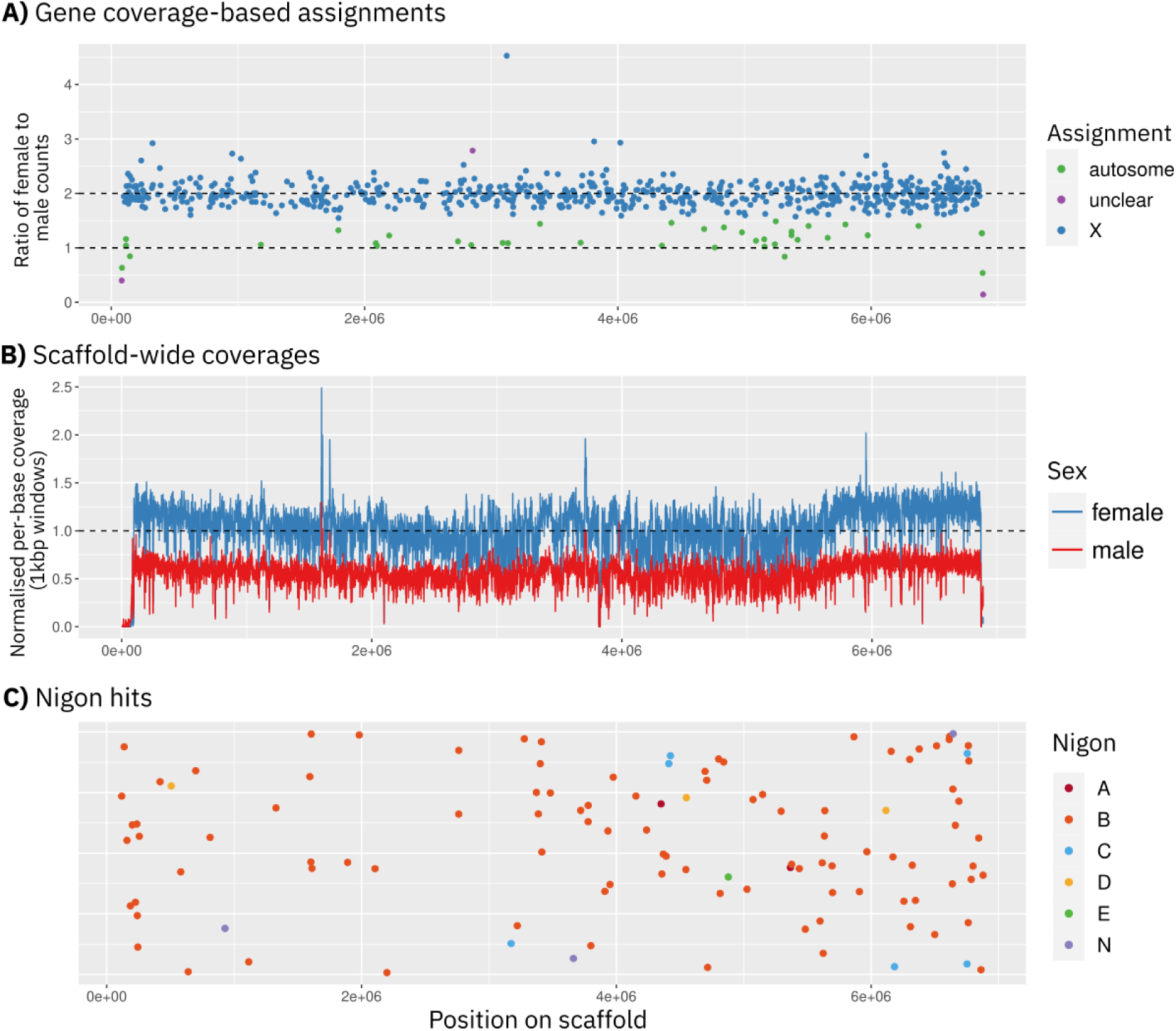
Example X-assigned scaffold in *M. monhystera* and its Nigon painting. **A)** Ratio of female to male DNA-Seq counts for genes on this specific scaffold: most genes have a ratio consistent with an X chromosome (blue dots). **B)** Normalised female and male DNA-Seq coverage along the entire scaffold (not just genes). Coverage was normalised so that diploid coverage should have a value of 1 (y-axis). Females have diploid coverage, and males haploid coverage, on this scaffold, again consistent with an X chromosome. **C)** Nigon painting of this X-assigned scaffold. Almost all Nigon elements along this scaffold are Nigon B assigned (orange dots). Y-axis values are meaningless here and used only for spreading out the points.

To infer the evolutionary origin of the sex chromosomes in our genome assemblies, we used Nigon elements. Nigon elements are the inferred ancestral chromosomes of nematodes in order Rhabditina (to which *Mesorhabditis* belongs), defined based on the observation that groups of genetic markers have mostly remained on the same chromosomes through evolutionary time. Previous work defined seven Nigon elements (A, B, C, D, E, N, X), with X being the ancestrally X-linked element (23,25), and assigned highly-conserved genes to each element (23). We thus ‘painted’ each of our confidently-assigned scaffolds by the Nigon-assigned genes found on it. As an example, one of the two X-assigned scaffolds of *M. monhystera* is mostly composed of Nigon B genes (Fig. 3C). We note that in the pseudosexuals, no Nigon elements were found on any Y-assigned scaffolds.

We initially expected to see many Nigon X hits across our X-assigned scaffolds, as its component genes were previously found on the X chromosomes of 15 nematode genomes (23); however, no *Mesorhabditis* genome was used in this definition set. Instead, we found that Nigon X elements were majoritarily on autosomes in all four *Mesorhabditis* species (68% for *M. spiculigera* and >90% for the other three *Mesorhabditis* species; Fig 4A and Fig. S1). Conversely, Nigon X elements were only partly present on the X chromosome of *M. spiculigera* (32%), and were near-absent from the X chromosome of the three other *Mesorhabditis* species (<3%).

**Fig. 4.**
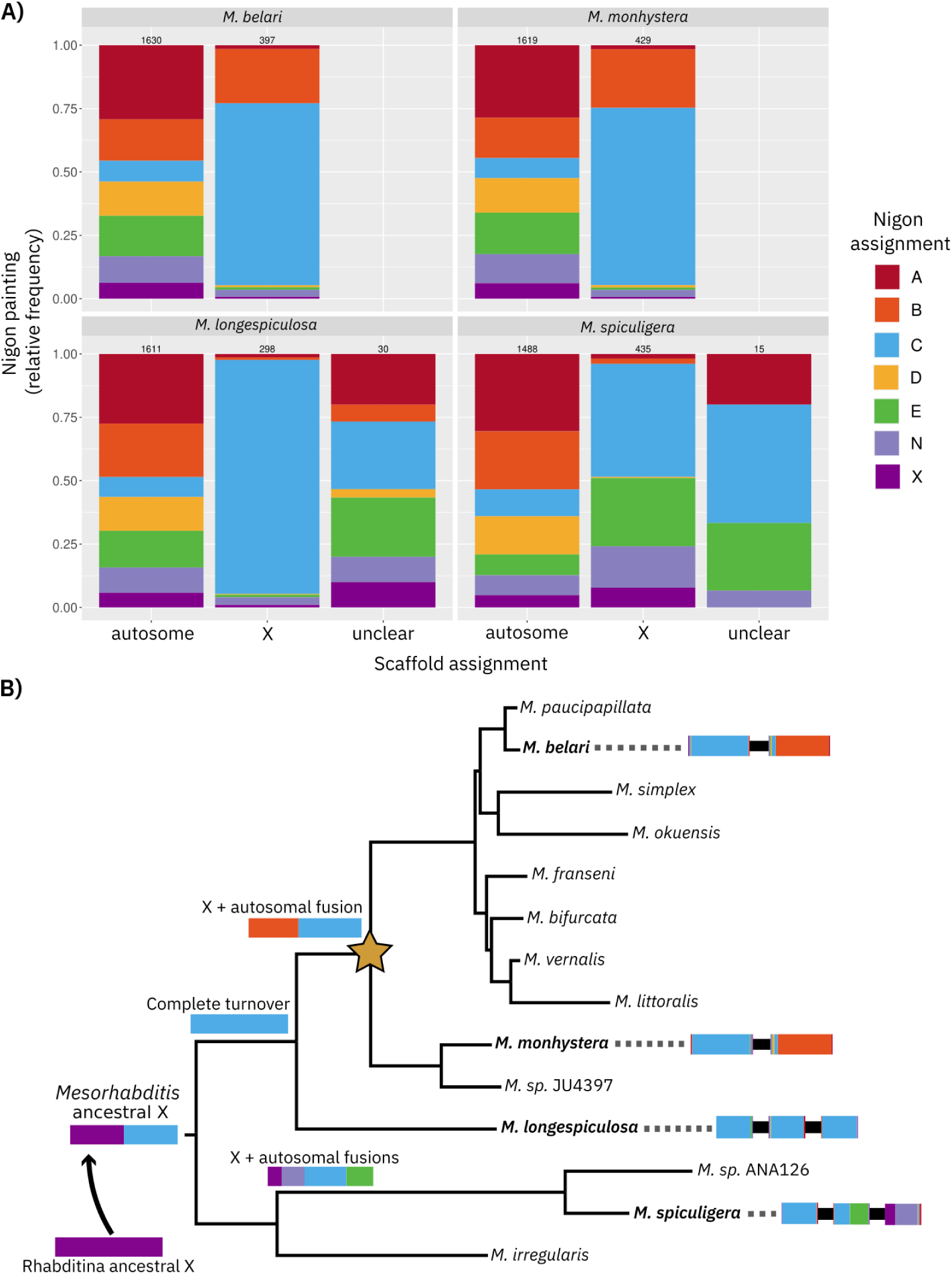
X chromosome turnover and fusions with autosomes in *Mesorhabditis*. **A)** For each scaffold assignment class (autosome, X and unclear, x-axis; see Methods for assignment criteria), the relative fraction of each Nigon element is shown as coloured bars (y-axis). Each panel shows one of the four focal species; numbers above coloured bars show the total number of Nigon elements per assignment and species. All X-assigned scaffolds have a significant fraction of Nigon C elements (blue), and the two pseudosexual species uniquely have Nigon B elements on their X chromosomes. **B)** The same tree as in Fig. 2 is shown, with a yellow star marking the origin of pseudosexuality. To the right of the tree leaves, putative X chromosomes are shown for our four focal species: each horizontal coloured bar represents one X-assigned scaffold, coloured by the relative distribution of Nigon elements on this scaffold, and black horizontal bars represent unbridged repeat arrays between scaffolds, preventing full assembly (see main text for details). To the left, at the tree nodes, the inferred ancestral state of the X chromosome at different points in time is shown. We infer an ancestral Nigon C fusion to the X at the base of all *Mesorhabditis*, as all four of our species have Nigon C painted X-assigned scaffolds. Fusion of Nigon E (green) and Nigon N (light purple) likely occurred in the lineage leading to *M. spiculigera*, and fusion of a Nigon B autosome (orange) occurred in the ancestor of all pseudosexuals. The Nigon B fusion co-occurs with the emergence of pseudosexuality.

Instead, we found the X chromosomes of all four species is majoritarily (54-70%) composed of Nigon C elements (blue, Fig. 4A). Assuming the ancestral X chromosome was composed of Nigon X (based on (23)), this clearly suggests the ancestral X chromosome of *Mesorhabditis* fully transitioned away into Nigon C, likely following a fusion event - except in the lineage leading to *M. spiculigera* (Fig. 4B).

For the two pseudosexuals, two X-assigned scaffolds were assembled in total, each mainly composed of Nigon B (orange) and Nigon C (blue) elements (Fig. 4A). To test whether these occur as two distinct physical units in cells, or as a single unit that was not contiguously assembled, we first used our Hi-C data. We found evidence for physical linkage in both species, but the overall signal intensity was relatively weak, especially in *M. belari* (see Methods and Tables S1/S2). We thus experimentally confirmed linkage of the two X-assigned scaffolds in *M. belari*, by genotyping animals across a pair of heterozygous loci (see Methods for details and Fig. S2). Our results strongly suggest Nigon B and Nigon C are physically fused in both pseudosexuals.

To test whether these fusion events were independent, we analysed the Nigon B elements on the fused autosomes. These were conserved and co-linear between *M. belari* and *M. monhystera* (Fig. S3), clearly supporting a single fusion event in the common ancestor of both species. Only part of Nigon B fused to the X chromosome in pseudosexuals, as the majority of Nigon B elements remain on autosomes in both *M. belari* and *M. monhystera* (Fig. S1). We thus conclude that, in the ancestor of pseudosexuals, part of an autosome (Nigon B) fused to the X chromosome (Nigon C; Fig. 4B).

We also found two additional fusions involving the X chromosome, both in the sexual species *M. spiculigera*: Nigon E (green) and Nigon N (light purple) (Fig. 4B and Fig. S4). For *M. longespiculosa*, all X-assigned scaffolds were Nigon C (blue).

Overall, the X chromosome has thus undergone fusions with various autosomes in different *Mesorhabditis* species, including a full replacement of the ancestral X chromosome by autosomal elements before the emergence of autopseudogamy.

### Emergence of the neo-Y chromosome in pseudosexuals

In an XX/X0 sex-determining system, X-autosome fusions can give rise to neo-Y chromosomes in males (Fig. 1B). While initially identical to the fused copy, if the unfused copy stops recombining in males it diverges from its autosomal counterpart (1), eventually showing haploid coverage in males.

In *M. belari*, the X-fused, Nigon B scaffold (orange in Fig. 4B) showed a pattern of male/female coverages exactly consistent with this scenario (Fig. 5A, upper panel). The right-hand part of this scaffold had the expected pattern for an X chromosome, with diploid coverage in females and haploid coverage in males, but the left-hand part was diploid in both males and females. This is compatible with the scenario shown in Fig. 1B, in which a neo-Y arises by fusion followed by divergence: here, one part of the Nigon B fusion still aligns with the X of females, making it diploid in males and corresponding to a pseudo-autosomal region, while the remaining part would have diverged, making it haploid in males. To test this scenario, we mapped Y-assigned scaffolds to the rest of the genome, and found that the best hits occurred at the junction of the diploid-haploid transition zone on this X scaffold (lower panel of Fig. 5A).

**Fig. 5.**
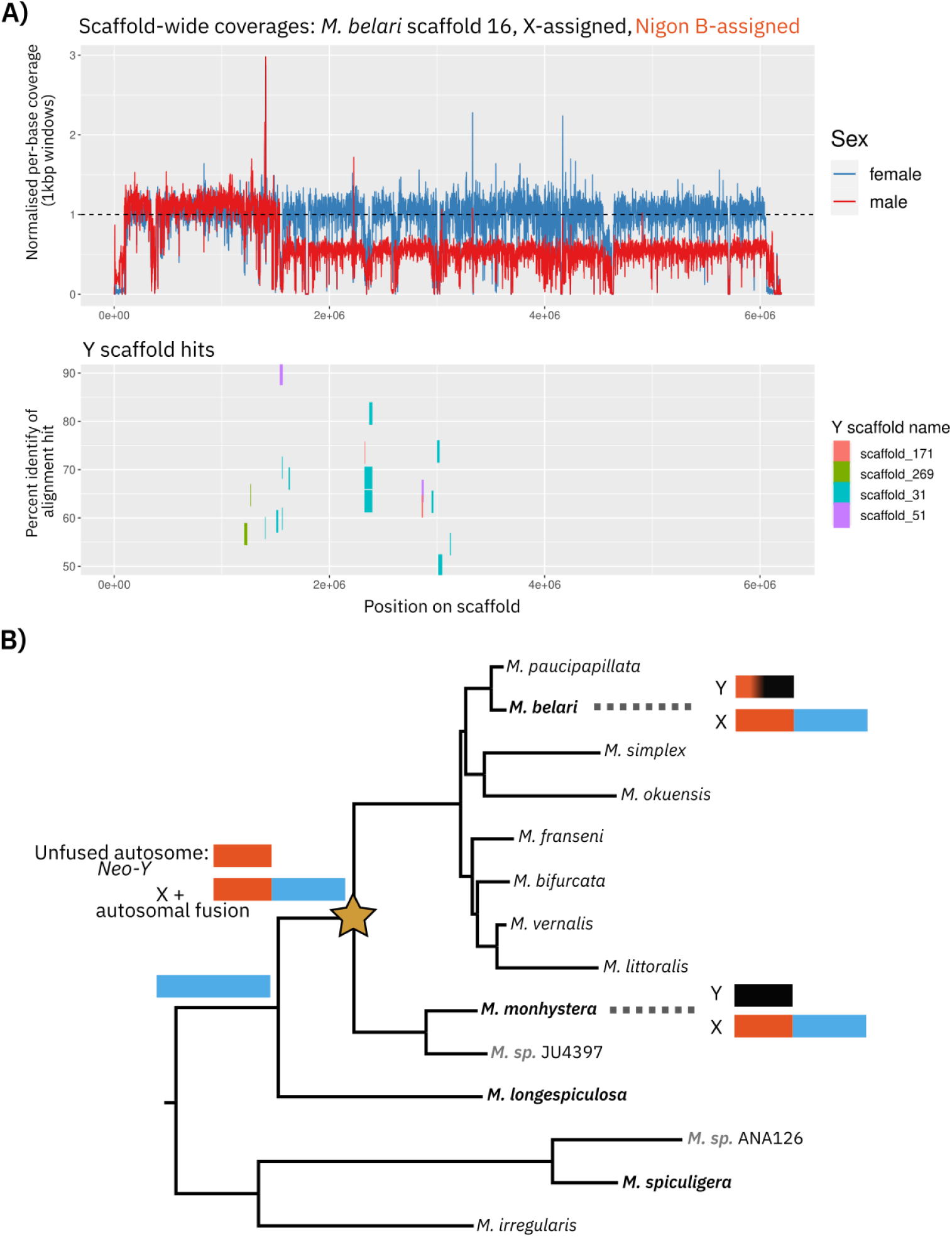
Neo-Y emergence following X-to-autosome fusion in the pseudosexuals. **A)** One of the two X-assigned scaffolds of *M. belari* (the one that fused to the ancestral X - see Fig. 4B) shows a mixture of autosome-like coverage (left-hand portion of the upper panel: females and males both have a coverage of 1, indicating diploid coverage) and X-like coverage (rest of the coverage plot, with diploid coverage in females and haploid coverage in males). To test whether the neo-Y chromosome of *M. belari* is related to this diploid region in males, we aligned Y-assigned scaffolds to the rest of the genome: the best hits genome-wide were found on this scaffold, and are displayed in the lower panel. The y-axis shows the percent identity of each alignment hit, the width of the bar represents how long the hit is, and the hit is coloured by which Y-assigned scaffold it was found in. The hits cluster in the transition zone between diploid and haploid coverage in males, consistent with the neo-Y originating from this entire scaffold. **B)** Evolutionary scenario for neo-Y origin in pseudosexuals. After the fusion of an ancestral autosome (orange) to the ancestral X (blue) in the lineage leading to pseudosexuals, the unfused autosome (orange) only ever occurs in males (assuming males remain defined by being haploid for the ancestral X, in blue). This neo-Y then diverged away from its autosomal ancestor: both are depicted as black rectangles, but they are of different sizes and do not share any large blocks of sequence homology (see Methods). Only in *M. belari* is a diploid region still found in males (shown as an orange portion on the neo-Y), suggesting recombination may still be occurring in this region.

These data are thus consistent with the neo-Y emerging following the Nigon B autosome fusion with the *Mesorhabditis* ancestral X chromosome (Fig. 5B). In *M. belari*, most of the neo-Y scaffolds did not map to the rest of the genome, showing it has substantially diverged in sequence from its fused counterpart in males - with the exception of the diploid region in males. Interestingly, in *M. monhystera*, no such diploid region was found, with the corresponding ‘B’ fusion being entirely haploid in males (Fig. 1B). This suggests the neo-Y of *M. monhystera* has fully diverged from its unfused counterpart, and possibly no longer recombines with the X.

We further tested this scenario using DNA stainings of meiotic cells in male gonads of the two pseudosexual species. In *M. monhystera*, we found two lagging chromosomes during metaphase and anaphase of meiosis I, suggesting they are not paired (Fig. S5). By contrast, in *M. belari* for which we expect proper pairing of sex chromosomes through the small pseudo-autosomal region (Fig. 5A), we did not observe any lagging chromosomes (Fig. S5). Lagging chromosomes were also absent in gonads of *M. monhystera* females, where the two X chromosomes are expected to pair (Fig. S5). This strongly suggests the unpaired chromosomes in *M. monhystera* males are the X and the Y.

### Characterisation of the neo-Y chromosomes

#### Overall neo-Y degeneration

We could confidently assign 20 scaffolds spanning 7.8Mbp to the Y chromosome of *M. belari*, and 152 scaffolds spanning 17Mbp to the Y chromosome of *M. monhystera*. Despite the large size of the neo-Ys, we found very few predicted genes with RNA-Seq expression: 155 for *M. belari* and 319 for *M. monhystera*. In both species, gene density (number of genes per kbp) was ∼8-10 fold lower on the Y than on X and autosomes. In both species, genes on the Y also had more paralogous copies, and were part of fewer Y-specific families, compared to genes on the X or autosomes (Tables S3 and S4). The Y chromosomes were also highly enriched for repeated elements compared to the X and autosomes (Table S5), and had a higher fraction of genes with no RNA-Seq support (see Methods). Altogether this is consistent with pseudogenisations, gene duplications and accumulation of repeated sequences on the neo-Y of pseudosexuals, as often seen during the evolution of Y chromosomes (3,4).

#### Age of the neo-Y

Based on a recent time-calibrated phylogeny of the clade Rhabditina (26), to which *Mesorhabditis* belongs, we estimate the neo-Y emerged ∼45 million years ago (see Methods). X–autosome fusions are frequent in some clades, yet XX/XO sex-chromosome systems persist, indicating that recently emerged neo-Y chromosomes following X–autosome fusions may easily be lost (as suggested in nematodes (20–23) and in insects (19)). We thus hypothesised that specific factors have contributed to maintaining the neo-Y in the pseudosexual *Mesorhabditis*.

#### No evidence for changes in genetic sex determination specific to the pseudosexuals

One reason for maintenance is if the Y is necessary for sex determination. In the XX/X0 nematode *C. elegans*, specific genes serve to count the ratio between X chromosomes and autosomes, leading to the specification of males or hermaphrodites (27,28). In *C. elegans*, these are *sex-1* and *fox-1* located on the X chromosome, and *sea-1* and *sea-2* located on autosomes (28). Together these control the dose of the ‘master’ sex regulator gene *xol-1* (28). As the two *Mesorhabditis* sexual species are XX/XO, they likely also rely on a X:autosome counting. We could not identify orthologs of *xol-1* and *sea-1*, perhaps in line with the fast divergence of *xol-1* already observed within the *Caenorhabditis* genus (29). However, we could identify orthologs for *sex-1*, *fox-1* and *sea-2* (see Methods and Supp. Data). All three orthologs - including the X-linked *sex-1* and *fox-1* in *C. elegans* - were located on autosomes; for *sex-1* and *fox-1*, this is likely due to the near complete transition of the X in *Mesorhabditis*. We also found *sex-1*, *fox-1* and *sea-2* in the pseudosexuals, also all located on autosomes. Importantly, the non-synonymous divergence dN of these genes relative to *C. elegans* was not higher in the pseudosexuals than in the closest sexual, *M. longespiculosa*, despite the synonymous divergence dS being saturated (>1; see Methods), indicating ample opportunity for degeneration. This suggests these three genes’ function has been preserved in the pseudosexuals, and thus that they may also rely on the same dosage mechanism for sex determination as their sexual counterparts - rather than on their Y chromosome.

#### Repeated Y fertilisation drive

In *M. belari*, Y-bearing sperm have a strong fertilisation drive. We previously found that of 30 amphimictic embryos (where the sperm DNA is used; Fig. 1A), all developed into males as adults, suggesting only Y-bearing sperm are competent for fertilisation. Next, using DNA-FISH with a Y-specific probe we found that ∼87% of all fertilised oocytes - including gynogenetic ones - are fertilised by Y-bearing sperm (18). Here we found that *M. monhystera* also has a fertilisation drive, using mounted one-cell embryos for DIC live imaging. Of 11 amphimictic embryos that developed into adults, 11 were males. At a p-value threshold of 0.05, this is incompatible with a fertilisation drive below 74% (chi-square goodness-of-fit test; see Methods). This result demonstrates that both *M. belari* and *M. monhystera* are subjected to a strong Y drive, suggesting that Y drivers may have contributed to the maintenance of the Y in the pseudosexual species.

To identify candidate drivers, we looked for protein-coding genes with annotations consistent with a role in the fertilisation drive. We first identified four families with particularly interesting annotations: putative transmembrane proteins, which could be cell-surface exposed in Y sperm only, and proteins with a putative role in meiosis, which could segregate proteins specifically to Y-bearing sperm. However, all had closely-related paralogs on the X or autosomes, making them unlikely candidates. We next identified genes occurring exclusively on the Y (i.e. with no paralogs on the X or autosomes): 91 genes in 10 families in *M. belari* and 63 genes in 17 families in *M. monhystera* (provided in Supp. Data). Of these, some also had putative function in a fertilisation drive, as DNA-binding proteins (helix-loop helix, zinc finger, TATA-box-binding domains), transmembrane proteins, and structural proteins involved in chromosome segregation (vimentin, condensin, kinetochore protein). However, these genes were not orthologous at the sequence level between the two pseudosexuals, i.e. were species-specific. Overall, we only identified species-specific candidate drivers on the neo-Y, suggesting driving factors either emerged after the speciation of *M. belari* and *M. monhystera*, or preceded their speciation but then diverged substantially.

The Y chromosome may not in itself encode a protein directly responsible for the fertilisation drive, however. Notably, we found - by detection of an antibody against the active form of RNA polymerase II - that in male gonads of *M. belari*, spermatocytes are transcriptionally silent from the onset of meiosis I, i.e., prior to segregation of the neo-Y (Fig. S6). This suggests that a factor responsible for the drive may not be expressed specifically in Y-bearing sperm after they have been individualised (though we note immunostaining may not mean that Y-bearing sperm are fully transcriptionally silent). Instead, factors responsible for the Y fertilisation drive, e.g a protein responsible for oocyte/sperm recognition, could be expressed earlier, and elsewhere (e.g. on the X or autosomes), and co-segregate with the Y during meiosis, perhaps through recognition of a non-coding element, such as a repeat array, on the Y.

To test this idea, we looked for repeated arrays of small (between ∼3bp and 120bp units) sequences called satellites, as these have been implicated in meiotic drives in *Drosophila* systems (30,31). We found several satellites highly specific to the Y chromosomes of both *M. belari* and *M. monhystera*, as well as a few satellites specific to their X chromosomes, but none of these satellites were in common between the two species (see Methods and Supp. Data).

Overall, we have thus identified a conserved fertilisation drive phenotype in the two pseudosexuals, but with no conserved candidate loci at the genetic level. We return to this in the Discussion.

#### Conserved male-beneficial gene families on the neo-Y

In addition to the fertilisation drive, which could have maintained the neo-Y since its emergence, we looked for any male-beneficial genes on the neo-Y that could also have played a role in its maintenance (32).

We first looked for any orthologous families with members on the Y chromosomes of both *M. belari* and *M. monhystera*, as such genes could have maintained the neo-Y prior to the split of the two species. Using whole sequence-based homology (with OrthoFinder (33)), we found just one such family that we called ‘*hen-1*-like’, as members are orthologous to the *C. elegans* gene *hen-1*. In *C. elegans hen-1* codes for an LDL receptor domain-containing secretory protein involved in sensory perception (34), and the members in *M. belari* and *M. monhystera* were similarly annotated as ‘lipoprotein receptors’ (using eggNOG (35)). In *M. belari*, the family had two members on autosomes and 15 on the Y, and in *M. monhystera*, one member on an autosome and 4 on the Y, consistent with a Y-specific expansion. In both species, the Y-specific copies bear specific amino acid substitutions compared to the autosomal copies that are highly conserved across the Y copies. The substitutions were specific to each species, however, suggesting the duplications on the Y may have occurred independently in the two species (i.e., after the split of *M. belari* and *M. monhystera*; Fig. S7).

Because the *hen-1*-like genes could be cell surface-exposed and/or secreted, we initially wondered if they were involved in oocyte-sperm interactions. However, comparing RNA-Seq data of gonads with whole adults in *M. belari*, we found that most of the expression was not in gonads (estimated 1-5% of the total expression in gonads; see Supp. Data), consistent with a function in somatic tissues, maybe for sensory perception as found in *C. elegans* (34). We thus hypothesise the *hen-1*-like genes on the Y chromosome encode male-beneficial functions, but are not involved in a fertilisation drive, as we expect such genes to be transcribed in the germline. We note that the autosomal copies already have a male expression bias, of 16% in *M. belari* up to 600% in *M. monhystera* (see Methods and Supp. Data).

As just one family of orthologs was common to the neo-Y chromosomes, we expanded our search to shared functional domains. We found only a few shared functional terms, of which ‘lipoprotein receptor’ - corresponding to the *hen-1*-like family - and ‘promoting spermatogenesis’. For the latter, we found two main gene families, one in each pseudosexual species, encoding zinc-binding transcription factors with homology to the *C. elegans* gene *tra-1*, a master regulator of germline and somatic sex differentiation (36,37). The homology to *tra-1* was particularly clear for the *M. monhystera* family (copies aligned well to *C. elegans tra-1*), and more distant for the *M. belari* family (only domain-level homology was found) (Fig. S8). In both families, 8-9 copies were found on the neo-Y chromosome, and 1-4 copies were found on the X chromosome, and the Y copies had species-specific substitutions, again consistent with male-specific evolution on the neo-Y (Fig. S8). Here also, the non-Y copies - located on the X chromosome - had a strong male expression bias, of 278% in *M. monhystera* and with almost zero female expression in *M. belari*.

While *tra-1* was initially identified as a strongly feminising gene in *C. elegans* (notably, XX mutants for *tra-1* become fertile males (38)), recent work has revealed a key role in spermatogenesis (36,37,39) as well as male behaviour, including exploration, through expression in specific neurons (40). We thus similarly hypothesise that the neo-Y *tra-1*-like copies encode male-beneficial functions.

Overall, we identified two putative male-beneficial gene families, *tra-1* and *hen-1*-like genes, on the neo-Y chromosomes of the two pseudosexual species, that could have contributed to its maintenance.

### Search for mitochondrial genome rearrangements

Finally, because the neo-Y chromosome is the only DNA that is transmitted exclusively through males, we also analysed the other DNA with uniparental transmission: the mitochondrial genome. We specifically wanted to test for genomic changes associated with the pseudosexuals - similarly to the neo-Y emergence. For this we assembled the mitochondrial genomes of all *Mesorhabditis* species shown in Fig. 2A using long-read data (mostly Nanopore; see Methods). The assembled mitochondrial genomes all contain 12 protein coding genes (*atp-8* being absent, as in most other nematodes (41)), and are relatively small, ranging 13-15kbp. We found two large-scale mitochondrial genome translocations along the phylogeny. The first is common to all pseudosexual species except *M. monhystera* and *M. sp. JU4397*, with the gene order of the latter being conserved with *C. elegans* (41). We also found a translocation in one of the sexual species, *M. irregularis*. We concluded that mitogenome translocations are frequent in this genera but are not linked to pseudosexuality (Fig. S9).

## Discussion

In this study, using genomics, we reconstituted the evolution of the sex chromosomes in the nematode genus *Mesorhabditis*. We discovered that a Y chromosome co-occurs with a reproductive system featuring a very low proportion of males whose genome is not transmitted to females, but only to sons (18). We found the closest extant sexual species in this genus are XX/XO - the predominant system in other nematodes. To our knowledge, this is the first evidence of a male-specific chromosome identified in animal species with rare males.

Unexpectedly, we found X chromosomes in *Mesorhabditis* that have entirely lost the features common to the X chromosome of other rhabditid nematodes (i.e., Nigon X); this could call for a redefinition of these common features in Rhabditina. Perhaps as a consequence of this evolutionary transition, we found that among well-known genes that count the X:autosome ratio in *C. elegans*, present on the X and on autosomes in *C. elegans*, all identified orthologs were found on autosomes in *Mesorhabditis*. Because we found no differences in rates of evolution of these genes between sexual (XX/X0) and pseudosexual (XX/XY) species, we hypothesise the Y chromosome of pseudosexual species does not play a role in sex determination. An interesting parallel is the genus *Drosophila* in which a Y chromosome is consistently found, but sex is determined using the dose of X-linked elements (42,43). However, how the X:autosome ratio is counted in *Mesorhabditis* remains to be elucidated.

Our data suggests a single X-autosome fusion gave rise to a neo-Y chromosome in pseudosexual species of *Mesorhabditis*. Since then, the neo-Y maintained a pseudo-autosomal region with the fused X in *M. belari*, but not in *M. monhystera*. These patterns are most likely driven by partial or full cessation of recombination between the two copies of the autosome portion in males. Cessation of recombination can occur directly at fusion points, as has been shown in *C. elegans* after induced chromosome fusions (44), or through subsequent inversions (45). Consistent with the former, the pseudo-autosomal region in *M. belari* is located distal to the fusion point, so that recombination may gradually have been lost from there. In *M. belari*, the pseudo-autosomal region could mediate recombination and proper chromosome segregation between the X and Y during male meiosis. In *M. monhystera*, we found no such pseudo-autosomal region, and how proper X/Y segregation is achieved in this species, likely in the absence of recombination, remains to be investigated.

Once recombination stops, degeneration of the neo-Y often occurs (1). Our identified neo-Y shows all the characteristic features of degeneration: accumulations of repeats, pseudogenisations and gene duplications (3). Given enough time and degeneration, neo-Ys can also be lost entirely (46). In rhabditid nematodes, the X chromosome has been especially prone to past fusions with autosomes (23) but most species are XX/X0, not XX/XY (24), so that any neo-Ys have mostly been lost. Here by contrast, we found a neo-Y that has been maintained for ∼45 millions of years in the pseudosexual species, suggesting factors conferring selective or transmission advantages in males have ensured its maintenance.

One possibility is the existence of male-beneficial genes on the neo-Y chromosomes. We identified two shared gene families on the Y chromosomes of the two pseudosexual species. One family is predicted to promote spermatogenesis (*tra-1*-like family) and the other to be involved in male learning and sensory perception (*hen-1*-like family). If truly male-beneficial, these could have maintained the neo-Y by conferring a selective advantage over males that have lost the neo-Y. Evolutionarily, both families underwent Y-specific duplications and substitutions that are specific to each species. These substitutions may thus have emerged only after the speciation of *M. belari* and *M. monhystera*; alternatively, they could have been fixed following gene conversions between the Y paralogs, a frequent phenomenon on Y chromosomes (47).

We also found that both Y chromosomes are involved in a strong oocyte fertilisation drive, which could provide a strong transmission advantage over males without a Y. We did not find any shared protein-coding genes or satellite repeats between the two pseudosexuals that could easily account for these drives. Evolutionarily, it could be that the Y driver loci are different between the two species, either because they have diverged in each species (e.g. through an arms race with the X chromosome (4)), or that drivers emerged independently in each species following neo-Y emergence. Another possibility is that long non-coding or small RNAs are involved, which was not explored in this study.

Mechanistically, we do not know whether a factor facilitates fertilisation by Y-bearing sperm or inhibits fertilisation by X-bearing sperm. Our observed drives could occur through expression of Y or X-bearing sperm following male meiosis (though they appear transcriptionally silent at this stage), or preferential segregation of a cytoplasmic factor to Y or X-bearing sperm during male meiosis. The latter has been observed before: in the nematode genus *Auanema*, proteins involved in sperm motility co-segregate with the X chromosome during male meiosis (48), and in *D. melanogaster*, the well-known driver Stellate poisons Y-bearing sperm during male meiosis (49).

In light of our results, we hypothesise that, in *Mesorhabditis*, the evolution of pseudosexuality and the Y chromosome may be causally linked. Indeed, because females make up ∼90% of populations, and because males do not contribute to the female genome, autosomes and the X chromosomes are under selection overwhelmingly in females. Any male-beneficial genes, ancestrally on X or autosomes, can thus rapidly disappear - unless they become encoded on a Y chromosome. It could thus be that neo-Y emergence preconditioned, or facilitated, the emergence of autopseudogamy. Once both a neo-Y and autopseudogamy occur, Y-chromosome drivers can then become rapidly fixed: as mostly gynogenetic eggs are produced that do not use the sperm DNA and give rise to females, Y-linked drivers do not strongly affect the sex ratio in the rare eggs that do use the sperm DNA, and can easily invade and become fixed. Conversely, a fertilisation drive of Y-bearing sperm ensures that when oocytes use the sperm DNA (in amphimictic embryos), males are mostly produced, and not sexual females - in turn, stabilising autopseudogamy.

In this study we demonstrated co-occurrence, but did not resolve the relative timing of autopseudogamy and neo-Y chromosome emergence. Future denser sampling could resolve this, by identifying sexual species with a neo-Y or autopseudogamous species without a neo-Y. Absence of both could further demonstrate a stabilising effect of the neo-Y on autopseudogamy. In addition, despite our reasoning and results, we cannot definitively exclude that the Y has no selected function. To truly test this, forward (unbiased) or reverse (starting with our candidates) genetic screens for Y genes affecting male fitness, and/or whose sperm exhibits more or less fertilisation bias, will be needed.

Overall, the rarity of male-specific chromosomes among known animals that have transitioned away from canonical sexuality suggests that reproductive systems involving rare males are more likely to emerge in lineages lacking differentiated male sex chromosomes - or, that any pre-existing Y chromosomes are easily lost. Our study illustrates that old male-specific chromosomes can in fact be maintained in species with rare males, and calls for a broader exploration of sex chromosomes in pseudosexual species.

## Materials and Methods

### Nematode culture and sequencing

All strains were maintained at 20°C on enriched plates (Nematode Growth Media supplemented with 7 g/L agarose, 5 g/L bacto-peptone and seeded with *E. coli* OP50). We sequenced the following *Mesorhabditis* strains: JU2817 (*M. belari*), JU2855 (*M. monhystera*), DF5017 (*M. longespiculosa*), AF72 (*M. spiculigera*), JU2848 (*M. littoralis*), JU2864 (*M. simplex*), JU3143 (*M. okuensis*), ANA126 (*M. sp.* ana126), JU3174 (*M. franseni*), JU3347 (*M. vernalis*), JU2902 (*M. bifurcata*), JU3003 (*M. paucipapillata*), JU4397 (*M. sp.* JU4397), SB321 (*M. irregularis*).

To produce worm pellets, worms were synchronised by bleaching as previously described in (18) and then grown as above. When plates were nearly starved, worms were harvested and washed in M9 + 0,01% Tween20. Worm pellets were stored at −20°C.

#### DNA Sequencing: M. belari and M. spiculigera

We performed high molecular weight DNA extraction of a 40 mg frozen pellet of *M. belari* using the Monarch® HMW DNA Extraction Kit for Tissue (NEB) with the following modifications. 50 µl of the lysis buffer mix was used for pestling the frozen worm pellet. We added the rest of the lysis buffer and transferred the mix in a 2 ml DNA LoBind^Ⓡ^ Tube (Eppendorf). The mix was digested at 56 ℃ in the Thermomix C (Eppendorf) at 750 rpm for 2 hrs. After RNA digestion we spun the sample for 5 min at 2500 rcf and transferred the supernatant to a new 1.5 ml DNA LoBind^Ⓡ^ Tube (Eppendorf) leaving about 30 µl behind. We added 30 µl of lysis buffer to the mix and put the sample for 5 min on ice. We added 300 µl cold Protein Separation solution and inverted the sample at about 20 rpm for 1 to 2 min. We put the sample for 10 min on ice. We spun the sample at 16.000 rcf for 25 min at room temperature. We transferred the supernatant to a new 2 ml DNA LoBind^Ⓡ^ Tube (Eppendorf) and added 2 glass beads. We added 550 µl Isopropanol and inverted the sample several times before we put it in a wheel for 5 min at 7 rpm. From here we followed the recommended protocol steps. In total, we extracted 2.640 ng high molecular weight DNA. 2,500 ng of this DNA was sheared in two rounds to an average size of 21.1 kb with a Megaruptor 3 (setting 28 followed by setting 29 in the second shearing round) (Diagenode). The sheared DNA was SPRI cleaned with 0.81x of ProNex beads (Promega).

For the DNA extraction of *M. spiculigera* we used CTAB digestion followed by Phenol-Chloroform extraction. We used a worm pellet of 50 mg for extraction. We pestled the frozen worm pellet in 300 µl CTAB lysis buffer (2% w/v) and added 10 µl Proteinase K (800 U/ml). After 1 hour digestion in a Thermomixer C (Eppendorf) at 56 ℃, 550 rpm, we added 10 µl RNaseA and left it for another hour in the Thermomixer. After a total of 2 hours we added 1x Phenol/Chloroform/Isoamylalcohol maix (25:24:1) and mixed well with a bore tip. We centrifuged at 14,000 rcf for 10 min at 4 ℃. We took the supernatant and performed two Chloroform washes with 1x Chloroform/Isoamylalcohol (49:1). We centrifuged 14,000 rcf for 10 min at 4 ℃. We then transferred the supernatant to a new 1.5 ml tube and added 0.7x Isopropanol. We inverted the mix and incubated it for 10 min at room temperature. We centrifuged at 14,000 rcf for 10 min at 4 ℃. We washed the pellet twice with 70% Ethanol and centrifuged for 5 min 14,000 at 4 ℃. The pellet was eluted in a volume of 200 µl elution buffer. We then mixed the extracted DNA several times with a bore pipette tip and let the DNA elute overnight at room temperature before proceeding to QC measurements. We extracted 4,500 ng DNA. We sheared the DNA to an average size of 21.4 kb with a Megaruptor 3 (setting 29) (Diagenode). The sheared DNA was SPRI cleaned with 0.81x of ProNex beads (Promega).

The PacBio libraries were prepared by the Sequencing Operations core at the Wellcome Sanger Institute.

Hi-C library preparation and sequencing were performed by the Scientific Operations: Sequencing Operations core at the Wellcome Sanger Institute. A 25 mg pellet of adult-enriched nematodes of *M. belari* strain JU2817 and *M. spiculigera* strain AF72 was processed using the Arima Hi-C version 2 kit following the manufacturer’s instructions. An Illumina library was prepared using the NEBNext Ultra II DNA Library Prep Kit and sequenced on one-eighth of a NovaSeq S4 lane using paired-end 150 bp sequencing.

#### DNA Sequencing: all other *Mesorhabditis* species

For DNA extraction, 100-200 μl of nematode pellets were snap frozen in liquid nitrogen and thawed at 37°C three times for breakage of the cuticle. These were vortexed in 600 μl of Cell Lysis Solution (Qiagen ref. 158906) and 6 μl of proteinase K (17 mg/ml; Sigma-Aldrich), then incubated three hours at 65°C with gentle shaking. After addition of 40 μl of RNAse A (5 mg/ml; Roche), samples were incubated 1 hr at 37°C, then 5 min on ice with 200 μl of Protein Precipitation Solution (Qiagen ref. 158910). After 10 min centrifugation at 13000 rpm, 4°C, the supernatant was precipitated with 600 μl of Isopropanol, incubated 10 minutes at room temperature and centrifuged 30 min at 13000 rpm, 4°C. The pellet was washed twice with 70% ethanol, centrifuged, dried 1h under a hood, then dissolved in 30 μl of water.

For sequencing, *M. littoralis*, *M. simplex*, *M. okuensis* and *M. sp.* ANA126 were processed by IGFL (Lyon, France), with library preparation using Native Barcoding Kit 24 V14 and sequencing on an ONT MinION MK1C machine (R10 flow cell).

*M. longespiculosa*, *M. franseni*, *M. vernalis*, *M. bifurcata*, *M. paucipapillata*, *M. sp.* JU4397 and *M. irregularis* were also processed by IGFL, with the same library preparation and sequencing on an ONT PromethION machine (R10 flow cell).

The *M. monhystera* sample was processed by BGI (China) for high-fidelity (HiFi) library preparation and sequencing on a PacBio Revio machine. In addition a worm pellet of *M. monhystera* was processed by the MGX platform (Montpellier France) for Hi-C library preparation (Arima-HiC kit) and paired-end sequencing (2×150bp) on an Illumina NovaSeq 6000 machine.

For *M. belari*, *M. monhystera*, *M. longespiculosa* and *M. spiculigera*, we also separately sequenced pools of male and female worms. *M. belari* and *M. spiculigera* samples were processed by the MGX platform with library preparation using Illumina’s DNA prep kit and paired-end sequencing (2×250bp) on an Illumina NovaSeq 6000 machine. *M. monhystera* and *M. longespiculosa* were processed by the GenomEast platform (France): *M. longespiculosa* was sequenced on an Illumina HiSeq 4000 (2×100bp) and *M. monhystera* on an Illumina NextSeq 2000 (2×100bp).

#### RNA Sequencing

For *M. belari* and *M. longespiculosa*, we prepared RNA from entire populations. 100 μl of nematode pellets were vortexed in 500 μl Trizol. The suspension was frozen in liquid nitrogen and transferred to 37°C water bath (4-5 times), then vortexed at room temperature for 30 s, let rest 30 s (5 times). 100 μl of chloroform (24:1 CHCl :isoamyl alcohol) was added, tubes were vigorously shaken for 15 s, incubated at room temperature for 2-3 min, then centrifuged 15 min at 13000 rpm, 4°C. The upper phase was transferred to a new tube with 250 μl of isopropanol. After mixing and incubation at room temperature for 10 min, the tube was centrifuged for 15 min at 13000 rpm, 4°C. The pellet was washed with 75% ethanol, then centrifuged for 5 min at 13000 rpm, 4°C. After removing the supernatant and the pellet was air-dried for 15-20 min, then dissolved with 50-100 μl of RNAse-free water.

For *M. belari*, *M. monhystera, M. longespiculosa* and *M. spiculigera*, we sequenced RNA from single adult males and adult females using the Smart-Seq2 protocol adapted for single worms and described in (50), with the following modifications: 1μl of proteinase K at 17mg/ml, SuperaseIn instead of RNAsin ribonuclease inhibitor from Promega, and an extra step of snap freezing in liquid nitrogen before worm lysis.

All samples were processed by the GenomEast platform, with library preparation using Nextera XT and paired-end sequencing (2×100bp) on an Illumina HiSeq 4000. For *M. monhystera* and *M. spiculigera*, the reads from males and females were pooled for gene prediction.

### Genome assembly

For three species (*M. belari*, *M. monhystera*, *M. spiculigera*) we had PacBio HiFi and Hi-C data for genome assembly. We first assembled the HiFi reads into contigs representing a single, unphased haplotype with hifiasm v0.19.5 (51) (`-s 0.55 --primary`). Haplotypic duplications (heterozygous regions collapsed into a single contig) were removed from the primary assembly with purge_dups v1.2.6 (https://github.com/dfguan/purge_dups). We then mapped the Hi-C reads to the purged assemblies with bwa v0.7.17 (52) (`bwa mem -SP5M`) and scaffolded the contigs with yahs v1.2a.2 (53). We then removed any contaminant scaffolds (e.g. bacteria coming from intestines of sequenced worms) with BlobToolKit v4.3.9 (54), based on abnormal coverage of the HiFi data or GC content and aberrant genes characteristic of non-nematode clades.

For the remaining species in the *Mesorhabditis* group (Fig. 2A), we had Nanopore data only. These were assembled with Flye v2.9.2 (55) (`-g 150m`), with two modes depending on how our Nanopore data was generated: `--nano-raw` for the MinION data and `--asm-coverage 50 --nano-hq` for the PromethION P2 data. For *M. longespiculosa*, we further purged and decontaminated the assembly as above.

For all assemblies, quality control was performed using BUSCO v5.8.0 (56) (with the genes in the ‘nematoda_odb10’ lineage) and Merqury v1.3 (57). For visual inspection of the scaffolding results, Hi-C contact maps were produced and visualised using Juicer (58). For metrics on the finished assemblies, see Table 1 below.

**Table 1.** Summary of assembly statistics and how much of each assembly was assigned to X/Y/autosomal/unclear status, for the four focal species of this study.

|  | *M. belari* | *M. monhystera* | *M. longespiculosa* | *M. spiculigera* |
| --- | --- | --- | --- | --- |
| Assembly span (#scaffolds) | 241 Mbp (454) | 211 Mbp (441) | 138 Mbp (725) | 147 Mbp (275) |
| Assembly N50 | 6.81 Mbp | 10.2 Mbp | 441 Kbp | 10.88 Mbp |
| %BUSCO genes found (%single-copy, %duplicated) | 83.6 (77.8, 5.8) | 84.1 (79.9, 4.2) | 83.1 (71.8, 11.3) | 81.1 (62.2, 16.9) |
| % autosome-assigned (span, #scaffolds) | 63.5 (153.2Mbp, 22) | 70.8 (149.4Mbp, 16) | 80.9 (111.5 Mbp, 394) | 70.7 (103.9 Mbp, 21) |
| %X-assigned (span, #scaffolds) | 6.3 (15.1Mbp, 2) | 8.1 (17.1Mbp, 2) | 8.32 (11.5Mbp, 20) | 15.4 (22.6Mbp, 6) |
| %Y-assigned (span, #scaffolds) | 3.2 (7.8Mbp, 20) | 8.1 (17.0Mbp, 152) | NA | NA |
| %unclear-assigned (span, #scaffolds) | 27.0 (65.3Mbp, 410) | 13.0 (27.4Mbp, 271) | 10.8 (14.9 Mbp, 311) | 13.9 (20.5 Mbp, 248) |

### Mesorhabditis phylogeny

After running BUSCO on each of our assembled genomes - plus *C. elegans*, used as an outgroup -, we found 422 genes present in single-copy across all assemblies (out of 3131 BUSCO searched genes). For each gene, we produced a multiple-sequence alignment of all protein-coding sequences with mafft v7.525 (59) and trimmed the alignment with clipkit v2.3.0 (60) to remove aligned positions with too many gaps (`-m smart-gap`). We concatenated all alignments into a supermatrix with seqkit v2.4.0 (61), from which we built a phylogeny using IQ-TREE v2.3.6 (62), with auto-detection of the best-fitting substitution model (here, VT+F+R7) and 1000 bootstrap replicates (`-B 1000`). Trees were visualised and rendered with FigTree (https://github.com/rambaut/figtree/).

### Gene prediction

We predicted protein-coding genes in the four focal species of this study (*M. belari*, *M. monhystera*, *M. longespiculosa* and *M. spiculigera*) using the tool BRAKER3 v3.0.7.5 (63). As input for prediction, we gave BRAKER3 a set of known proteins: the 3131 proteins in BUSCO ‘nematoda_odb10’, the proteins in BUSCO ‘metazoa_odb10’, and the existing proteomes of *C. elegans* and *M. belari* obtained from WormBase ParaSite release 18 (WBPS18) (64). We also gave BRAKER3 RNA-Seq sequencing data we obtained for each species: population-wide for *M. belari* and *M. longespiculosa*, and of adult males and females for *M. monhystera* and *M. spiculigera*. The RNA-Seq reads were mapped to our assembled genomes using STAR v2.7.10b (65) prior to use with BRAKER3. This resulted in 16717 predicted genes for *M. spiculigera*, 19410 for *M. longespiculosa*, 26653 for *M. belari* and 22542 for *M. monhystera*.

### Assigning genes and scaffolds to X, Y, or autosomal status

To assign individual genes a status we used our male and female DNA-Seq data to count the number of sequencing reads coming from each gene with kallisto v0.50.1 (66). For comparability between males and females, we used the size factor normalisation in DESeq2 (67). We assigned a status to each gene by taking the closest expected regression line of male (*m*) vs female (*f*) counts for X, Y or autosomal genes. In other words, we assigned each gene to the status minimising the following squared errors *e*:

- Autosomal gene: expectation: *f* = *m*, *e* = (*f* - *m*)^2^
- X gene: expectation *f* = 2*m*, *e* = (*f* - 2*m*)^2^
- Y gene: expectation *f* = 0, squared error *e* = *f* ^2^

Genes with low read counts are hard to distinguish this way (Fig. 2B). We thus reassigned genes to a fourth status, ‘unclear’, if they had fewer than 30 read count support in either males (for X, Y, or autosomal genes) or females (for X or autosomal genes). To ensure confident assignment of Y genes, we also set any Y-assigned gene with >5 read count support in females to unclear.

To assign entire assembled scaffolds to X, Y or autosomal status, we used two approaches. In the first, we reused our assignments of individual genes found along each scaffold: a scaffold was assigned to a given status if >50% of its component genes were assigned to that status (including ‘unclear’), and to ‘unclear’ if no gene assignment group matched this criterion, or the scaffold had no genes.

In the second approach, we used coverages of male and female DNA-Seq reads computed along the entire scaffolds. Reads were mapped to assembled genomes with minimap2 v2.26 (68) (-x sr for short-read preset), duplicates marked with GATK’s picard v4.3.0 (69). To ensure coverage was computed only for unique and well-mapped regions of the genome, reads with MAPQ<20 (corresponding to a >=1% probability of the read being mapped to the wrong location, as estimated by minimap2) or marked as ‘SECONDARY’, ‘SUPPLEMENTARY’ or ‘DUP’ were filtered out with samtools v1.17 (70). We computed coverage along 1kbp non-overlapping windows of scaffolds with bedtools v2.31.0 (71), and normalised coverages so that a value of 1 corresponded to the expected coverage on a diploid chromosome. For this, we divided the value in each window by the median coverage along all 1kbp windows in scaffolds of size >1Mbp, assuming that most of the genome is composed of autosomes. We used large scaffolds only for the median as we observed these mostly had uniform coverage along their entire length. For assignment, we define the median normalised female coverage as *n_f_* and that for males *n_m_*, and the normalised female/male ratio *r* = *n_f_* / *n_m_*. We then used the following ad-hoc rules:

- Autosomal scaffold: 0.6 < *n_f_* < 1.4 and 0.6 < *n_m_* < 1.4 and 0.7 < *r* < 1.3
- X scaffold: 0.6 < *n_f_* < 1.4 and 1.5 < *r* < 2.5
- Y scaffold: 0.05 < *n_m_* and *n_f_*< 0.001
- Unclear scaffold: neither of the above

Note that for Y assignment, a value close to 0.5 is expected in males, as they should be haploid in males. However, due to their high repeat content, most coverage windows had very low coverage values, as reads did not map confidently to a single location. When this constraint was lifted (i.e., no longer filtering reads by MAPQ), a value of ∼0.5 was restored in males: an example is shown in Fig. S10. We filtered including MAPQ, and with a low minimum coverage in males (0.05), to ensure that almost no coverage in uniquely mappable regions was found in females (0.001 threshold above).

We then proceeded to an inspection of these assignments to produce a final classification. If both gene and scaffold coverage-based classification were in agreement, we used this classification (1397 out of 1895 scaffolds). In all remaining cases, either the gene-based or the coverage-based classification were ‘unclear’. In most cases we set the scaffold to ‘unclear’ classification (379 of the 498 disagreeing scaffolds set to unclear), especially when few genes were on that scaffold. We note that many of these 379 scaffolds are in fact eliminated portions of the genome, due to Programmed DNA Elimination in *Mesorhabditis* nematodes (72): their normalised coverages in males and females were typically <0.6 because eliminated portions of chromosomes are only sequenced in germline cells, which form a minority of sequenced individuals. For the remaining scaffolds, we nonetheless assigned eliminated scaffolds if they contained many genes, to ensure accurate Nigon painting-based analyses, and to ensure we captured all genes on the Y chromosomes. All our classifications, and the data supporting them, are publicly available: see data availability section.

### Summary of genome assemblies and scaffold assignments

In Table 1 below, we summarise overall assembly and scaffold assignment statistics.

We note that the main reason our assemblies are still highly fragmented, and that many small scaffolds were of unclear assignment in all four species, is because of the process of Programmed DNA Elimination (PDE) that occurs in the genus *Mesorhabditis* (72). During PDE, somatic cells undergo the systematic destruction of parts of their genome during early embryonic development. As mixed-stage, whole individuals were sequenced here, both somatic and germline cells are captured and the genome assembly contains a mixture of eliminated (=present only in germline cells) and retained (=present in both somatic and germline cells) DNA. Most of the unclear-assigned scaffolds are eliminated: they have lower coverage and are highly enriched for long arrays of repetitive DNA, making contiguous assembly challenging. We defer a more detailed explanation of how we classified scaffolds as retained or eliminated to a future study on PDE in *Mesorhabditis*. However, these scaffolds have almost no BUSCO hits, and we manually checked that they did not correspond to X or Y scaffolds so as not to miss any possibly important genes, especially on the Y.

We also predicted the fraction of X, Y and autosomal-assigned scaffolds covered by repeated elements, using EarlGrey v3.1 (73): these data are shown in Table S5.

### Nigon painting

To paint our scaffolds with Nigon elements, we used the approach defined in (23), in which a subset of 2298 genes out of the 3131 BUSCO genes that are part of the ‘nematoda_odb10’ lineage were each assigned a Nigon letter (A, B, C, D, E, N, X). This letter was determined on the basis of gene co-occurrence patterns in chromosome-level assemblies of 12 species in the order Rhabditina (23). The correspondence table between BUSCO gene ID and Nigon letter is publicly available here: https://github.com/tolkit/nemADSQ/blob/main/example/gene2Nigon_busco20200927.tsv.gz. For visualisation, we used the same colour code for Nigon letters as in (23).

### Testing for X chromosome linkage

#### Testing for X linkage using Hi-C

To test whether the two X-assigned scaffolds of *M. belari* and *M. monhystera* were physically linked, we mapped the Hi-C sequencing data to our assembled genomes using bwa v0.7.17 (`bwa mem -SP5M`) and extracted contacts using two approaches. The first and most permissive is using pairix v0.3.9 (`bam2pairs`) (74), as all aligned reads are used to compute contacts, including reads aligning to repetitive parts of the genome. The second is using pairtools v1.1.2 (75), that parses contacts in a more sophisticated manner and allows filtering aligned reads by MAPQ, a score proportional to the probability that a read is misaligned (essentially due to sequence repetition in the genome). We computed contacts using pairix, and pairtools for reads with minimum MAPQ of 1 and 20.

To test for linkage, we then extracted all contacts overlapping the first or last 50kbp of each assembled scaffold and counted the number of contacts supporting each possible pair of scaffolds. For a given scaffold pair, four possible extremity contacts are possible, based on whether the contacts occur at the start or end (i.e., 5’ or 3’ end) of each scaffold. For each species, we extracted the top 10 pairs by inter-scaffold count, normalised by how much contact occurs at the extremities themselves (=within-contact count), and looked for whether the two X-assigned scaffolds consistently came on top. This was clearly the case for *M. monhystera*, even after filtering for reads with MAPQ>=20 (Tables S1 and S2).

#### Testing for X linkage by PCR-based genotyping

To further test for physical linkage in *M. belari*, we used a PCR-based approach. We first identified heterozygous polymorphisms on each of the two X-assigned scaffolds (scaffold 7 and scaffold 16) whose size difference can be easily distinguished on agarose gels. We targeted variants at the 3’ end of these scaffolds as based on the Hi-C data, this is the likely position of these scaffolds’ linkage. Using IGV (76) we found a heterozygous 182-bp insertion on scaffold 7 (at position 8,594,943, in the intron of gene g23849), and a heterozygous 124-bp insertion on scaffold 16 (at position 5,122,107, in the intron of gene g8032). We designed PCR primers amplifying both the short and long alleles, with expected amplicon sizes of 395bp and 577bp for scaffold 7 (forward primer 5’-TCACCCGTTCTTCCCATTACA-3’, reverse primer 5’-TGGCTCGGTAGGGGAAATTT-3’) and 437bp and 541bp for scaffold 16 (forward primer 5’-TGTTTCCCATGTAAGTTGAGCA-3’, reverse primer 5’-TCTTCTTCAGGACCGATCTCC-3’). We also designed primers to amplify a homozygous and unique region of the neo-Y (on scaffold 31, of size 436bp; forward primer 5’-TGGCTTTTAGATCAATGGAACCT-3’, reverse primer 5’-GCAACGGTGATTTTCAAGCA-3’), to confirm genotyped individuals were male. We confirmed using PCR on wild-type individual *M. belari* males and females (strain JU2817) that the X-linked markers were indeed heterozygous in females, and haploid in males with the short or long allele, and that only males were positive for the Y marker (data not shown, see results below).

We then isolated a single adult gravid female on an agar plate (containing nematode growth medium and seeded with *E. coli* strain OP50), and lysed 27 individual male descendents of this initial female (mostly second-generation males, as a given female only produces ∼5 males across her lifespan), plus one second-generation female as a control (lysis: 10μl worm lysis buffer + 1% proteinase K for each worm). We genotyped all these individuals for the two X and one neo-Y markers by PCR.

The results are shown in Fig. S2. The female amplified both X markers at the expected size and not the Y marker, while males amplified the Y marker, and one of the X markers, at the expected sizes. A total of 6 males were discarded for analysis as at least one of the X loci did not amplify strongly, leaving 21 males. Of these, and denoting ‘S_7_L_16_’ to mean ‘short allele for scaffold 7 and long allele for scaffold 16’, we obtained the following marker combination counts: 10 S_7_L_16_, 11 L_7_S_16_, 0 S_7_S_16_, 0 L_7_L_16_.

Under the null hypothesis that these two markers are on different chromosomes, all four combinations should occur at the ∼25% frequency due to Mendelian segregation at meiosis. By contrast if the markers are on the same chromosome, two of these combinations should be highly over-represented among males due to genetic linkage, and the other two combinations arise only after maternal X chromosome recombination exactly between the two markers. Under the null, the probability of observing data at least as extreme as ours (p-value) is 10e-04 (goodness-of-fit test on a one-dimensional contingency table, df = 3). We thus concluded that these two markers are on the same chromosome, and that a single X chromosome occurs in germline cells of *M. belari*. We note that for 4 of the 6 discarded males, the faint bands also supported the S_7_L_16_ or L_7_S_16_ combinations as found above; one male had two bands for the scaffold 7 locus; and the remaining male did not amplify any locus (Fig. S2).

### Assessing homology between neo-Ys and the rest of the genome

All Y-assigned scaffolds were extracted and aligned to the rest of the genome using minimap2 v2.26 (68) (`-x map-ont`). The resulting alignment files (PAF format) were processed in R and alignment blocks filtered for a MAPQ>20, number of aligned bases >5000 nucleotides and a percent identity > 50%. For the *M. belari* Y chromosome, the largest summed alignment blocks totalled 422 kbp against an X-assigned scaffold (scaffold_16; shown in Fig. 5B). The next longest alignments spanned 235kbp on an autosome (scaffold_20), but they were almost all composed of a single repeated element on scaffold_20 shared across many Y-assigned scaffolds. For *M. monhystera*, the largest summed alignment blocks totalled only 29kbp on an autosome (scaffold_11).

We also aligned the Y-assigned scaffolds between the two pseudosexual species, also using minimap2. Applying the same filters, no alignment blocks were found.

### Search for sex determining genes

We identified the well-known X:autosome ratio counting genes from *C. elegans* based on (28): the two X-linked counters *sex-1* and *fox-1*, the two autosomal counters *sea-1* and *sea-2* and the integrator *xol-1*. These were extracted from the *C. elegans* proteome downloaded from Wormbase Parasite Release 18, and the longest isoform of each protein sequence searched against each of our four *Mesorhabditis* genome assemblies using miniprot v0.13-r248 (77). No hits were found for *sea-1* or *xol-1*. For the remaining genes, a single hit was found across all four species: all hits were on autosomes, and consistent in terms of Nigon elements. In particular, *sex-1* and *fox-1*, both found on the X chromosome of *C. elegans*, were found in *Mesorhabditis* on Nigon X for *fox-1* and Nigon N for *sex-1*. These two Nigon elements compose the X chromosome of *C. elegans* (23) but have become mostly autosomal in *Mesorhabditis* (see main results).

To assess conservation levels, the coding sequences of the hits found in the *Mesorhabditis* species, and the *C. elegans* representative, were aligned at the codon level, using seaview v5.1 (78). The alignments were loaded into MEGA v12.1 (79) and synonymous (dS) and non-synonymous (dN) pairwise divergences between each *Mesorhabditis* hit and the *C. elegans* sequence were computed using the Nei-Gojobori method with Jukes-Cantor correction for multiple substitutions. Only positions with no alignment gaps in any sequence were used. The dS values were well above 1 in all cases and could thus not be reliably estimated. The dN values ranged 0.56-0.61 for *fox-1*, 0.81-0.93 for *sex-1* and 0.96-1.33 for *sea-2*, and were highly similar across the sexual *M. longespiculosa* and the two pseudosexual species, *M. belari* and *M. monhystera*. The dN value of the sexual species *M. spiculigera* was clearly higher than the other three species for *sea-2*, but not *fox-1* and *sex-1*.

The data tables of alignment hits on the genome, coding sequence alignments and dN values are provided as Supp. Data on Zenodo (see data availability section).

### Age estimation of the ancestral neo-Y

Our results support neo-Y emergence following a single X-autosome fusion event in the ancestor of the two pseudosexual species *M. belari* and *M. monhystera* (Fig. 4B). As a conservative estimate, we thus used the speciation point between *M. belari* and *M. monhystera* in our *Mesorhabditis* phylogeny (Fig. 2A) as the latest possible time of origin of the neo-Y.

We then used a recently time-calibrated phylogeny of the nematode clade Rhabditina (also called clade V), to which *Me*s*orhabditis* belongs (26). Their calibration relied on mutation rates and generation times as estimated in the genus *Caenorhabditis*. In their phylogeny (Fig. S3 of (26)), the two species *M. belari* and *M. spiculigera* diverged ∼86 million years ago. From this, we compared the point of speciation of *M. belari* and *M. monhystera* with that of *M. belari* and *M. spiculigera*, that forms the leftmost point in our *Mesorhabditis* phylogeny (Fig. 2A). We obtained two different age estimates, depending on whether we used *M. belari* or *M. spiculigera* as the reference for the present - e.g., due to different substitution rates in the two species - : 39.7 million years using *M. belari* and 51.45 million years using *M. spiculigera*. We thus estimated the age of the neo-Y at ∼45 million years.

### Testing for a fertilisation drive in *M. monhystera*

To test for a fertilisation drive, we isolated young gravid females, dissected their embryos, and mounted them on agarose pads for live DIC imaging. We looked for one-cell embryos containing a decondensed male pronucleus, indicative of an amphimictic embryo, before placing these on individual enriched plates, and identifying the sex of the resulting adult individual phenotypically. Of 16 amphimictic embryos, 5 did not hatch, and 11 hatched to become adult males. We then performed a chi-square goodness-of-fit test in R, testing the null hypothesis that viable amphimictic embryos were fertilised by X- and Y-bearing sperm at 50% frequency, in which case males and females are also expected at 50% frequency. At a p-value threshold of 0.05, observing 11/11 males is incompatible with a Y-bearing fertilisation rate below 74%.

### Analysis of genes on the neo-Y chromosomes

To exclude pseudogenes, we only analysed genes with some evidence of expression in either adult males or adult females. We used three adult male and two adult female samples for *M. belari* and *M. monhystera*, and one male and one female sample for *M. longespiculosa* and *M. spiculigera* (fewer biological replicates as they do not have any Y genes). Reads were trimmed using fastp v0.23.2 (80) (`--cut_mean_quality 20 --cut_by_quality3 --qualified_quality_phred 15 --length_required 50`) and transcript counts obtained using kallisto v0.50.1 (66), mapped against all predicted coding sequences (see ‘Gene prediction’ section above). Counts were normalised using the size-factor normalisation function of DESeq2 (67) in R.

We then filtered out genes with less than a mean normalised RNA-Seq count of 10 in either males or females. For *M. belari*, this removed 6.2%, 16.8% and 36.7% of all X, autosomal and Y genes respectively, and for *M. monhystera* this removed 5.8%, 11.2% and 28.2% of all X, autosomal and Y genes respectively. We note that Y-assigned genes also more frequently had no RNA-Seq support at all: ∼7% of all genes for both species, compared to 0.4-1.5% for X and autosomal genes. In *M. belari*, 9 Y-assigned genes had significant RNA-Seq support in females and were discarded from analysis.

#### Search for male-beneficial or candidate fertilisation driver genes

To analyse patterns of orthology and paralogy of genes on the neo-Y chromosomes, we first predicted families of homologs using OrthoFinder v2.5.5 (33). We used the predicted protein sequences of our four focal *Mesorhabditis* species, plus the proteome of *C. elegans* obtained from WormBase ParaSite WBPS18 (64) to identify any known *C. elegans* homologs for genes of interest. OrthoFinder groups genes into families called ‘Orthogroups’ that include both orthologs (homologs across species), and paralogs (homologs in the same species, i.e. arising from a gene duplication). We then looked for Orthogroups with members on the neo-Y of both pseudosexuals (*M. belari* and *M. monhystera*) and Orthogroups with members exclusively on the neo-Y of either pseudosexual (i.e., no paralogs on autosomes or the X chromosome).

To functionally annotate genes, we used eggNOG v2.1.12 (35) on the proteomes of our four focal species, and also used a table of known functions of *C. elegans* genes, parsed from the WormBase ParaSite proteome file, for when a *C. elegans* ortholog was found with OrthoFinder.

To analyse the evolution of specific gene families, we used the same methods as for the *Mesorhabditis* phylogeny (see above): alignments were produced using mafft, trimmed using clipkit, and trees produced using IQ-TREE. Alignments were visualised using seaview version 5.1 (78).

We used orthology to *C. elegans* and domain-level annotations from eggNOG to look for species-specific genes that could be candidates for the fertilisation drive of the neo-Y chromosomes. In *M. belari*, we found three different families, encoding *smc-5* genes (involved in meiosis), condensins (involved in chromosome compaction), and *bub-1* genes (a kinetochore protein, involved in chromosome segregation), all of which could be linked to meiosis, and thus to biased segregation of proteins to Y-bearing sperm. In *M. monhystera*, we found one family encoding PRG-1 genes, part of the Argonaute family, which has been tied to a meiotic driver in *D. melanogaster* (81). However, all of these had closely-related paralogs on the X or autosomes, bearing only a few Y-specific substitutions, making them unlikely candidates.

As most genes occurring exclusively on the neo-Y of either pseudosexual had no annotations from eggNOG, we further ran hh-suite v3.3.0 (82) (`hhblits -d uniclust30 -p 90 -noaddfilter -e 0.0000001`) on each of these genes to obtain further domain-level annotations. We then filtered out any hits with a homology probability <90%, E-value >1, or with no associated functional terms.

All alignment files and annotation tables of our candidates of interest are provided as Supp. Data on Zenodo (see data availability section).

### Analysis of satellite repeats

We used two approaches to identify satellite sequences. In the first, we used Tandem Repeats Finder (trf) v4.10.0-rc.2 (83) (`2 7 7 80 10 50 2000 -h -ngs`) to identify satellites in the Y- and X-assigned assembled scaffolds of *M. belari* and *M. monhystera*. We filtered the raw outputs of trf to include only satellites with a minimum size of 5bp, tandem copy number of 10, and minimum identity to the consensus repeat of 80%. Our candidate satellites were then searched in the entire assemblies using seqkit v2.4.0 (`seqkit locate`), and satellites with a minimum total copy number of 40, and with at least 70% total occurrences on Y- or X-assigned scaffolds (compared to the rest of the genome) were extracted. This yielded 32 Y-enriched and 5 X-enriched satellites in *M. belari* and 112 Y-enriched satellites in *M. monhystera*. None of the satellites were shared between species.

Because satellite repeats are hard to assemble, we also used k-seek (84) to discover satellites from sequencing reads directly. Detection was performed on our male and female Illumina DNA-Seq reads (used for assigning scaffolds, see above) for *M. belari* and *M. monhystera*. All satellite counts were normalised by the median coverage of the reads along our genome assemblies, so that counts correspond to the number of satellites per 1x sequenced position. We then looked for satellites with a count of at least 40 in males and with >4 times more counts in males compared to females. This yielded 9 Y-enriched and 8 Y-enriched satellites in *M. belari* and *M. monhystera*, respectively (after excluding 5 Y-enriched satellites in *M. belari* already found using the previous approach). Again, none of these were shared between species. To be less stringent, we also found no common satellite with >2 times the counts in males, and at least 10 copies.

All the species-specific satellites we identified across both approaches are provided as Supp. Data on Zenodo (see data availability section).

### Analysis of mitochondrial genomes

#### Mitochondrial genome assembly

To reconstruct the mitochondrial genomes, we re-used the DNA-Seq data for the same *Mesorhabditis* species that we used to build our phylogeny (Fig. 2A). Reads were cleaned using fastp (80). The genomes of all species were assembled using Flye (55). Mitochondrial genomes were then assembled and annotated using MitoHiFi (85) with the default annotation tool (86) and using the *C. elegans* mitochondrial genome as reference (NC_001328.1). To ensure absence of contamination, mitochondrial genome coverage was measured. For this, the contig identified as the mitochondrial sequence in the Flye assemblies was removed and replaced by the assembled mitochondrial genome from MitoHiFi, as the former tends to contain multiple copies of the mitochondrial genome (due to its circularity). Reads were mapped against the assembly using minimap2 (68) and the coverage estimated using samtools (70).

#### Mitochondrial phylogeny

To reconstruct the mitochondrial phylogeny, we used the annotated genes from MitoHiFi. For each species, 12 mitochondrial protein coding genes were aligned with PRANK (87) using a codon model and sites ambiguously aligned were removed with Gblocks (88). The 13 genes were concatenated and and a phylogenetic tree was built using PhyML v3.0 (89) under a GTR+G+I model with 100 bootstrap replicates and was rooted using the clade [*M. sp* ana0126*, M. irregularis and M. spiculigera*].

### Reproducibility

Analyses were performed using Nextflow workflows (90), with individual analysis steps run using Charliecloud containers (91) or Conda environments with built tools available on Bioconda (92). All used tools are thus publicly available via Bioconda. The workflows and their configuration and analysis code supporting this work are publicly available at https://doi.org/10.5281/zenodo.15407995.

## Data Availability

All assembled genomes and raw sequencing data are available at the ENA under project accession xx.

Data tables underlying the analyses and figures in the paper are publicly available at https://doi.org/10.5281/zenodo.15407995.

## Author Contributions

Conceptualisation: BL, MD

Data Curation: BL

Funding acquisition: MD

Investigation: BL, LS, NS, AH, EW, MD

Methodology: BL, LS, NS, MD

Software: BL

Visualisation: BL, NS

Resources: MK, CL, MB

Writing - original draft: BL, MD

Writing - review & editing: BL, LS, NS, MB, MD

## Acknowledgements

For computing, we gratefully acknowledge the resources of the HPC at ENS de Lyon (managed by the Pôle Scientifique de Modélisation Numérique (93)) on which all workflows were deployed and run.

This work was supported by a grant from the Agence Nationale de la Recherche to MD: ANR-19-CE02-0012.

We also thank Quentin Helleu and Cécile Courret for productive discussions.

