## Supplementary Materials for "Emergence and stabilisation of a neo-Y chromosome in nematode species with rare males"

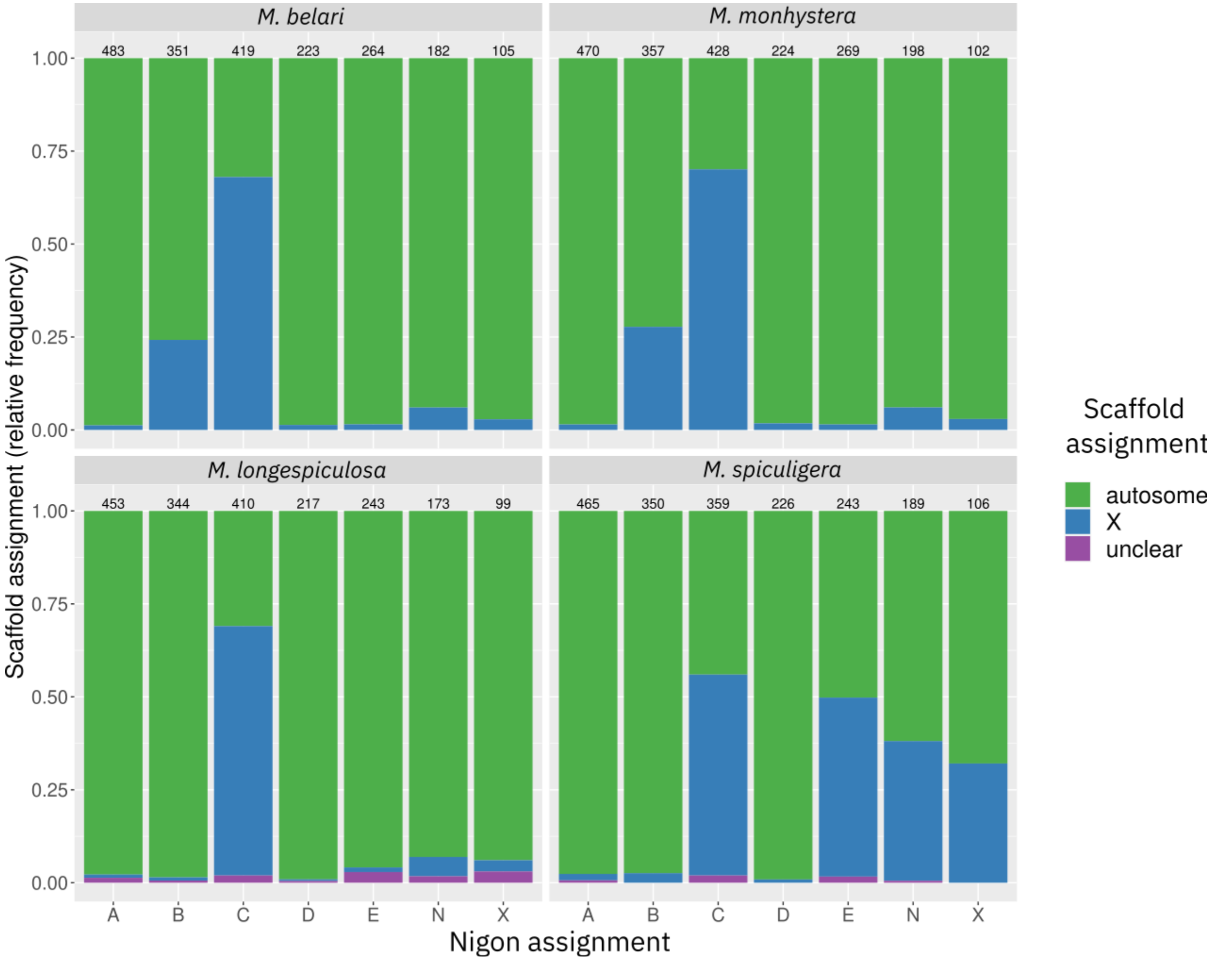


**Fig. S1. Average X/autosomal/unclear assignments for each Nigon and each of the four studied species.** For each species (panels) and each Nigon (x-axis), the relative distribution of that Nigon's presence on X, autosome - or 'unclear' if the scaffold's assignment was not clear-cut (see Methods) is shown. This allows assessing whether a given Nigon is more or less fully assigned to the X or to autosomes. For all four species, a majority of Nigon C is assigned to the X chromosome. Only part of Nigon B is on the X chromosome in the two pseudo-sexual species (lower panels); the same holds for Nigons N, X and E for *M. spiculigera* (upper-left panel). Nigon X, the ancestral X in the order Rhabditina (to which *Mesorhabditis* belongs), has become fully autosomal in all but one species here (*M. spiculigera*).

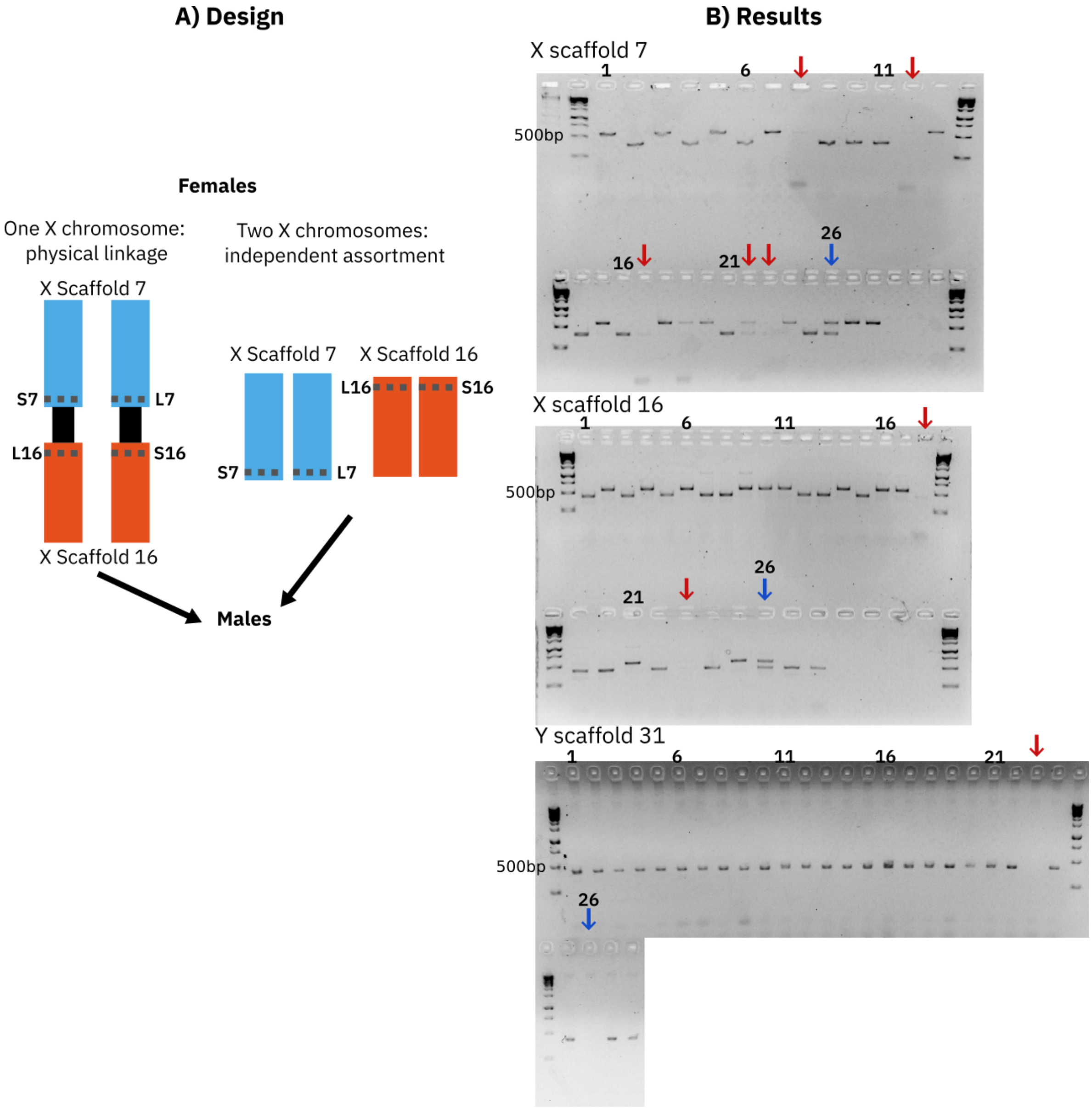


**Fig. S2**. **PCR-based validation of the X-autosome fusion in *M. belari*. A) Experimental design.** In *M. belari*, two X-assigned scaffolds were identified, scaffold 7 and scaffold 16, coloured here by their majority Nigon assignment (see main text). In females, where two X chromosomes occur, either these two scaffolds are physically linked (left-hand part), or they occur as distinct units (right-hand part, i.e. two X chromosomes in cells). To distinguish these, we identified heterozygous indels on each scaffold, near their extremities, shown as dotted lines. 'S7' means 'short allele on scaffold 7', 'L7' means 'long allele on scaffold 7', and similarly for scaffold 16. We then genotyped males descended from a single female, in which a single copy of each X chromosome is expected. If the two scaffolds are distinct units, the four possible combinations of the two indels should occur at similar frequencies (25%), due to independent assortment under Mendelian segregation. If the two scaffolds are physically linked, two of the four combinations should be disproportionately observed, the other two being produced by recombination events. **B) Results.** 27 males and one female were lysed and PCR amplified for each X chromosome indel, plus a Y chromosome locus (on scaffold 31, see Methods). Numbers above wells refer to genotyped individuals. The female is shown with a blue arrow (well number 26), and amplifies two bands at the X loci - confirming the locus is heterozygous - but does not amplify the Y locus, as expected. For the remaining wells, males amplified a single band at the Y marker, confirming they were male, and a single band at the same size as one of the two female bands for the two X markers. The expected sizes are: S7, 395bp; L7, 577bp; S16, 437bp; L16, 541bp;

Y scaffold 31, 436bp. To the left, the ladder band at size 500bp is shown. For wells marked with red arrows, bands were too faint and those individuals were discarded from the analysis. For the remaining 21 males, the following combinations of X genotypes were observed: 10 $S_7L_{16}$, 11 $L_7S_{16}$, 0 $S_7S_{16}$, 0 $L_7L_{16}$. Thus physical linkage is clearly supported; see Methods for statistical support.

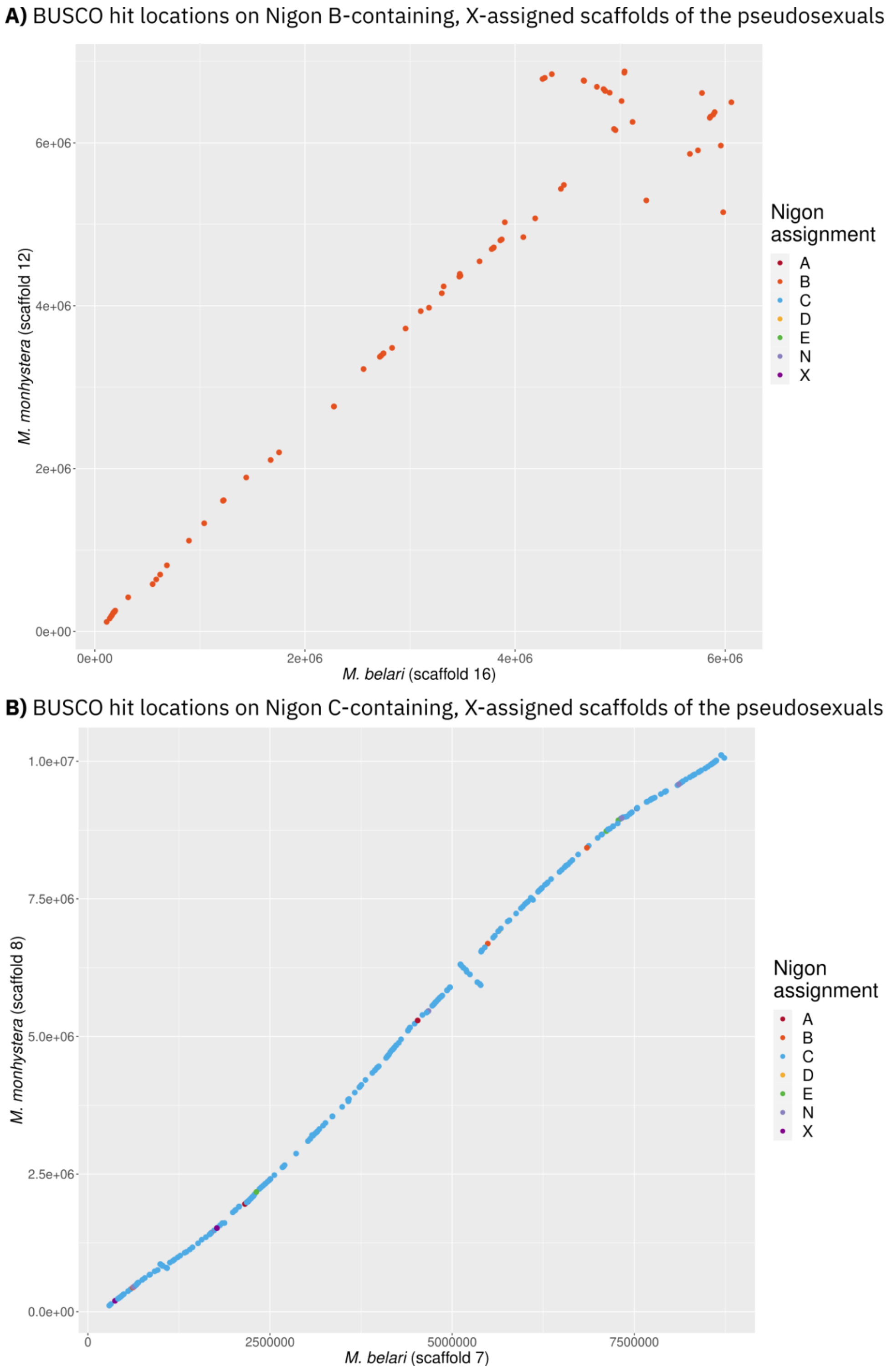


**Fig. S3. Common evolutionary origin of the X-assigned scaffolds in the two pseudosexuals.** Two X-assigned scaffolds were found for the two pseudosexual species: scaffold 12 and scaffold 8 in *M. monhystera*, and scaffold 16 and scaffold 7 in *M. belari* (Fig. 4B). To test for common evolutionary origins of these scaffolds across species, we compared the pairs with the same Nigon paintings: scaffold 12 and scaffold 16 (Panel A, Nigon C; orange) and scaffold 8 and scaffold 7 (Panel B, Nigon B; blue). In each panel, we plot the occurrence of each Nigon element on each scaffold. In both comparisons, the elements largely occur in the same relative order (except for one inversion in each), clearly supporting a common evolutionary origin.

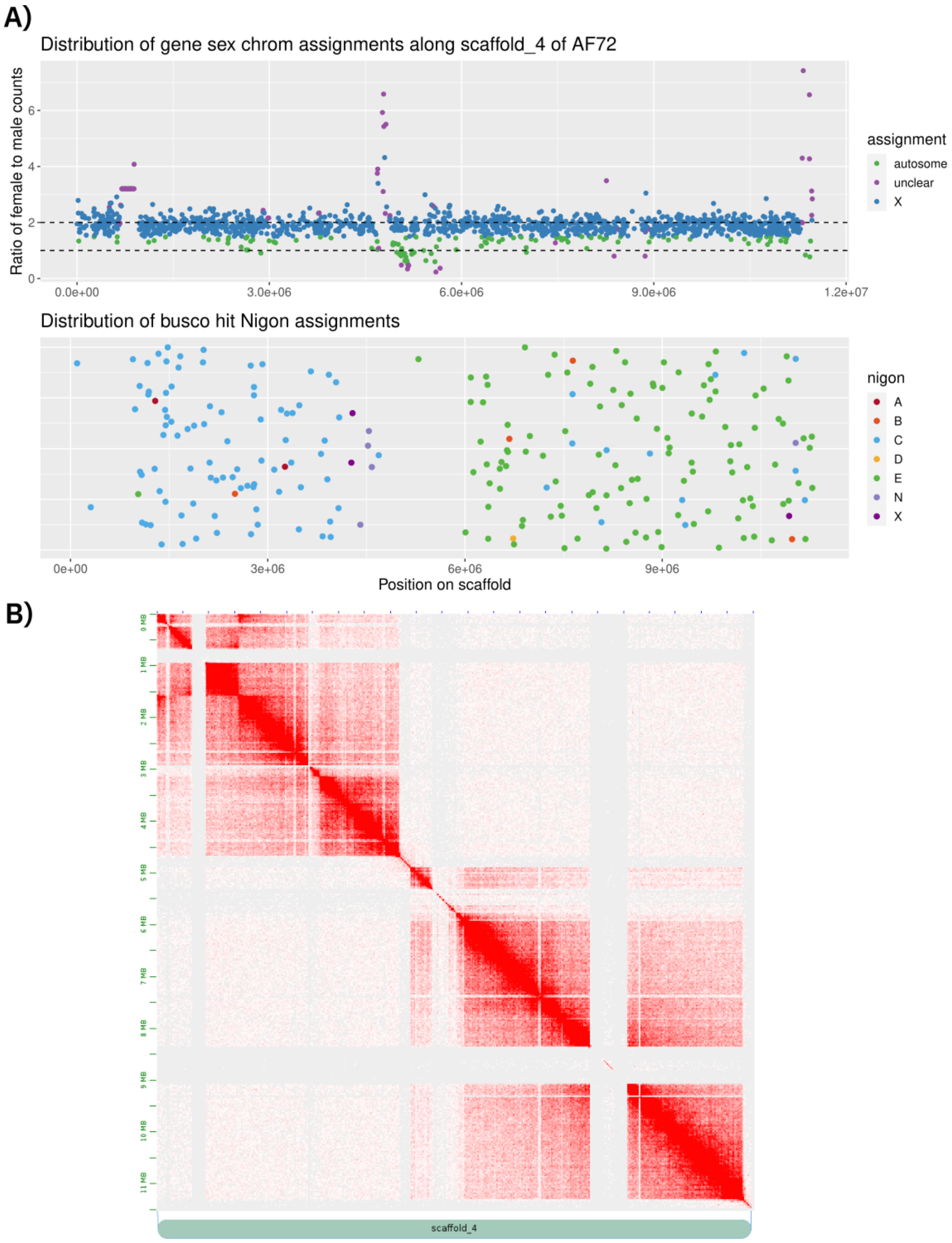


**Fig. S4. X-to-autosome fusion in *M. spiculigera*. A)** The scaffold is X-assigned (upper panel), with a majority of genes showing twice as much normalised coverage in females compared to males (blue dots in upper panel), and is a mixture of Nigons C and E (lower panel). **B)** The scaffold's Hi-C map, with red dots showing contacts along the scaffold's length. Portions with lower density of red dots, on the diagonal, correspond to repeated sequences. Despite this, the map is consistent with a single, well-assembled physical unit, supporting a true fusion event.

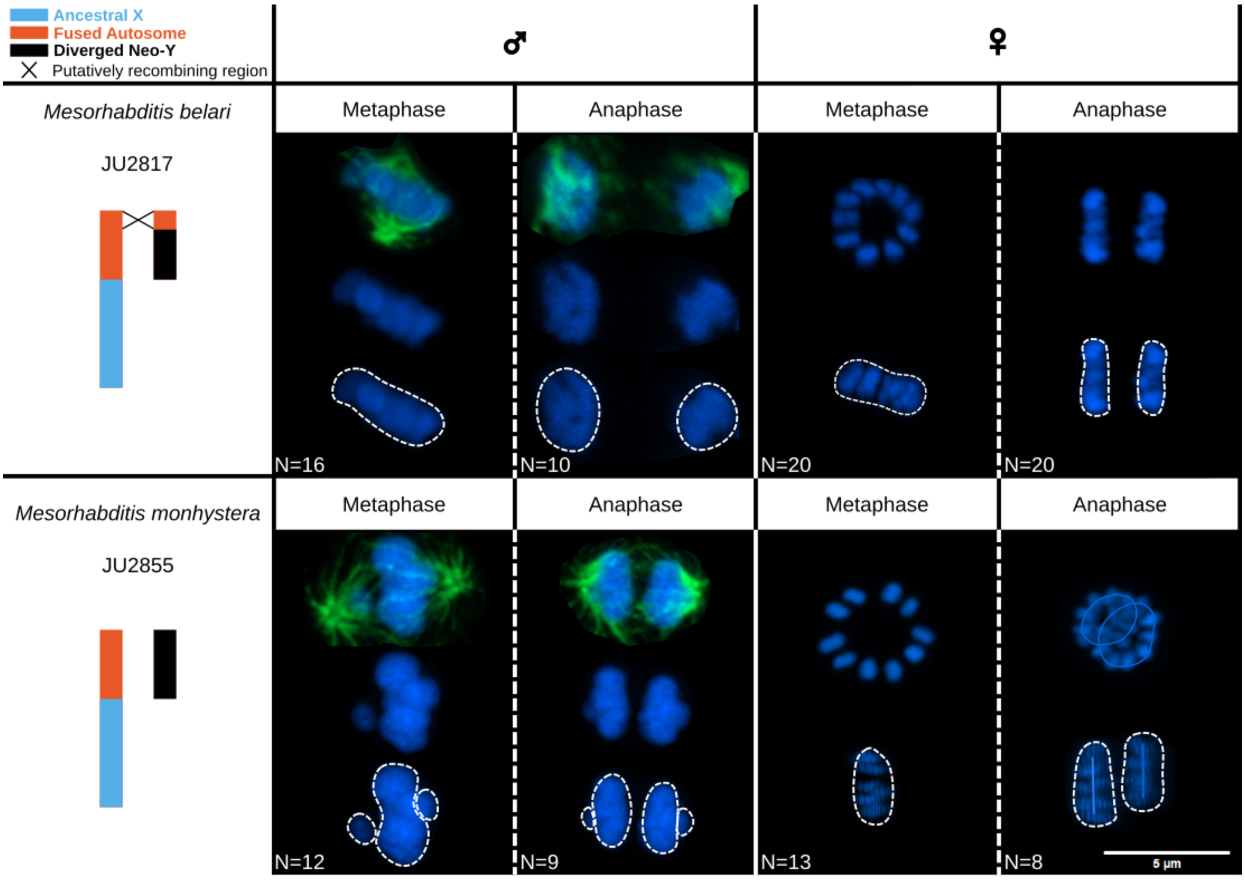


**Fig. S5. Two lagging chromosomes observed during male meiosis I in *M. monhystera*.** On the left, the X and neo-Y chromosomes are depicted, as in Fig. 5B. The diploid region in *M. belari* is shown, that putatively still recombines. On the right, representative cells in metaphase or anaphase of meiosis I are shown, from *M. belari* (upper panels) and *M. monhystera* (lower panels), in male (left-hand panels) and female (right-hand panels) gonads. DNA is in blue (Hoechst) and tubulin in green (immunostaining using an antibody against alpha-tubulin). White dotted lines show manual delimitation of the DNA content. In male gonads of *M. belari* and female gonads of both *M. belari* and *M. monhystera*, a single mass of DNA is found in metaphase I, and two masses of DNA in anaphase I, as expected if all chromosomes are properly paired. In both species, a characteristic 'rosette' of 10 sets of paired chromosomes can be seen in female metaphase I. By contrast, in male gonads of *M. monhystera*, two additional smaller masses of DNA can be found, in both metaphase I and anaphase I. Because these were never found in males of *M. belari*, nor in females of *M. monhystera* - whose only difference is possessing two X chromosomes that should be paired - we propose these are the X and Y chromosomes of *M. monhystera*, and that the X and Y are unpaired during male meiosis I in this species. In each panel, the number of observed cells (N=) that showed this pattern is provided.

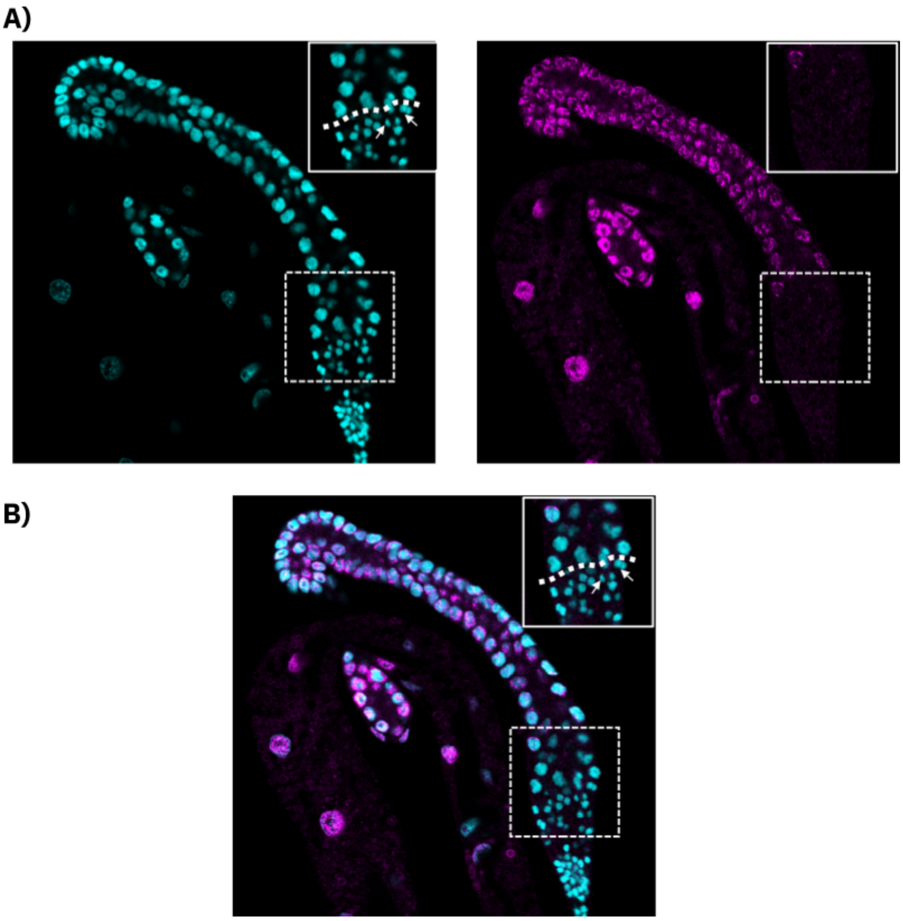


**Fig. S6: Spermatocytes are transcriptionally silent from the first meiotic division. A)** Gonad of a *M. belari* male, with DNA in cyan blue (left-hand panel) and active transcription in magenta (right-hand panel), as measured using immunostaining with an antibody against the phosphorylated form of Serine 10 on RNA Polymerase II. The gonad is spatially organised: the distal part, top left, contains the spermatocytes in meiotic prophase, which progressively enter the division phases. The proximal part (bottom right) contains the mature spermatids recognisable by their compact DNA. The dividing sperm cells are shown with the white arrows (metaphase and anaphase figures) and the transition is delineated with a white dotted line in the insets. The RNA Pol II signal is no longer detectable from the onset of the meiotic divisions. **B)** Merge of the DNA and RNA Pol II channels.

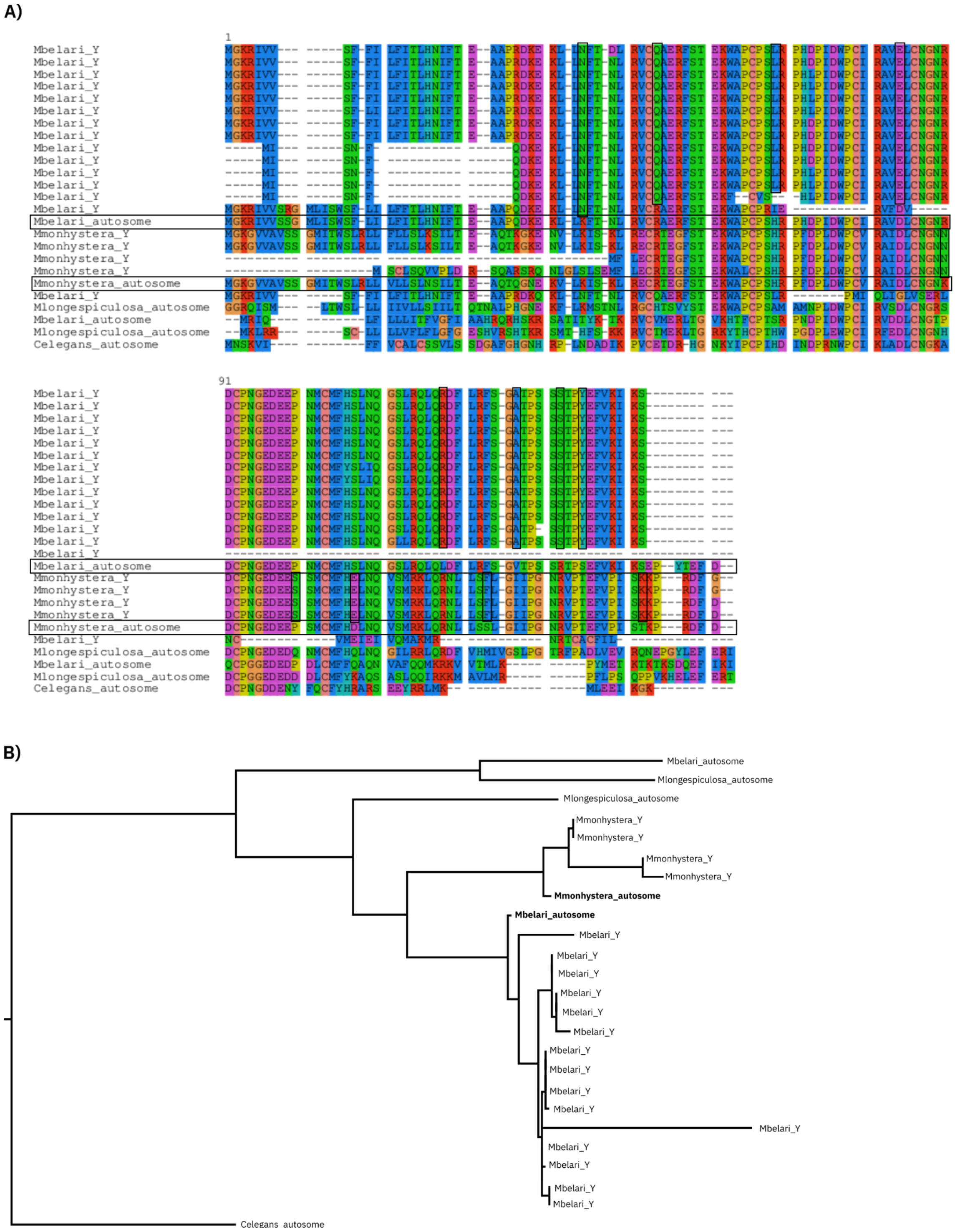


**Fig. S7: Evolution of the hen-1-like family of orthologs. A)** Multiple sequence alignment of the members of the hen-1-like family. On the left, the species in which the gene was found, and the chromosome it was found on, are indicated. In *M. belari*, 14 Y copies and two autosomal copies are found, and in *M. monhystera*, 4 Y copies and one autosomal copy was found. The *C. elegans* protein *hen-1* is shown in the last row of the alignment. The horizontal boxes highlight the autosomal copies from which the Y copies derived, and the vertical boxes highlight amino-acid substitutions that occurred specifically on the Y copies compared to the autosomal ancestor. These substitutions are specific to each of the two species. **B)** Phylogenetic tree built from the multiple sequence alignment, with the autosomal ancestor for *M. belari* and *M. monhystera* marked in bold.

A)

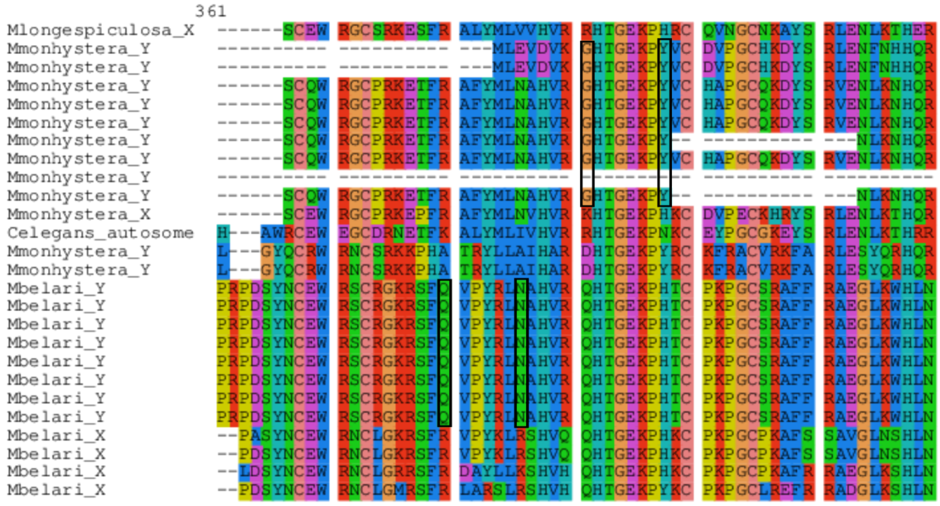


B)

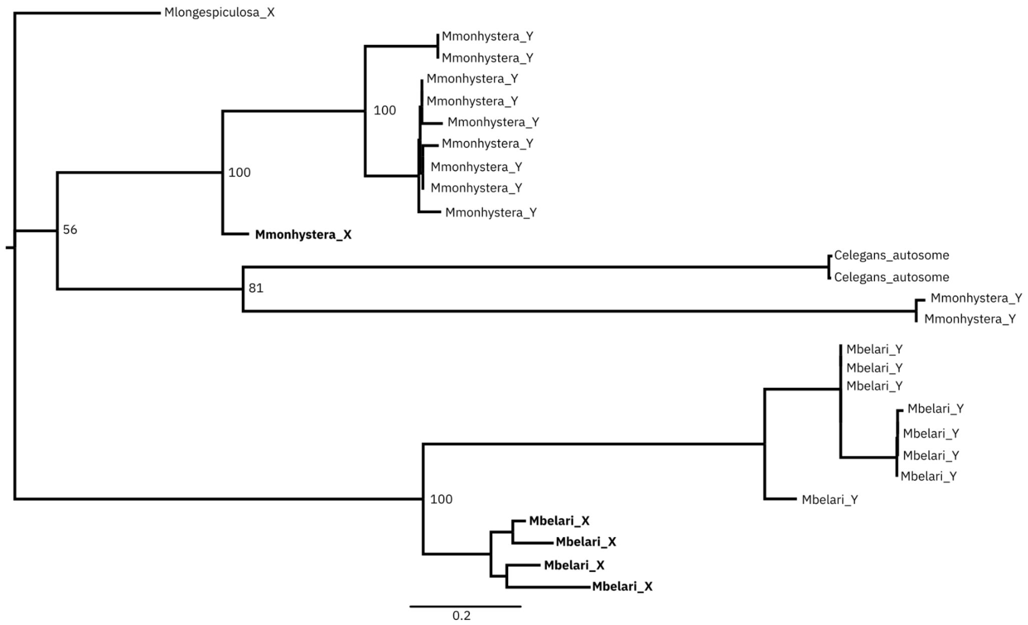


**Fig. S8: Evolution of the *tra-1*-like family of orthologs. A)** One region of the multiple sequence alignment of the members of the tra-1-like family is shown. The *C. elegans* copy is autosomal, while the *M. monhystera* and *M. belari* copies are on the X and neo-Y chromosomes. The vertical rectangles mark positions where the copies on the Y have acquired specific substitutions compared to the copies on the X chromosome. These substitutions are species-specific. **B)** Phylogenetic tree built from the full multiple sequence alignment, with the copies on the X marked in bold. Numbers on nodes indicate bootstrap support and scale bar (bottom) is in amino acid substitutions per site.

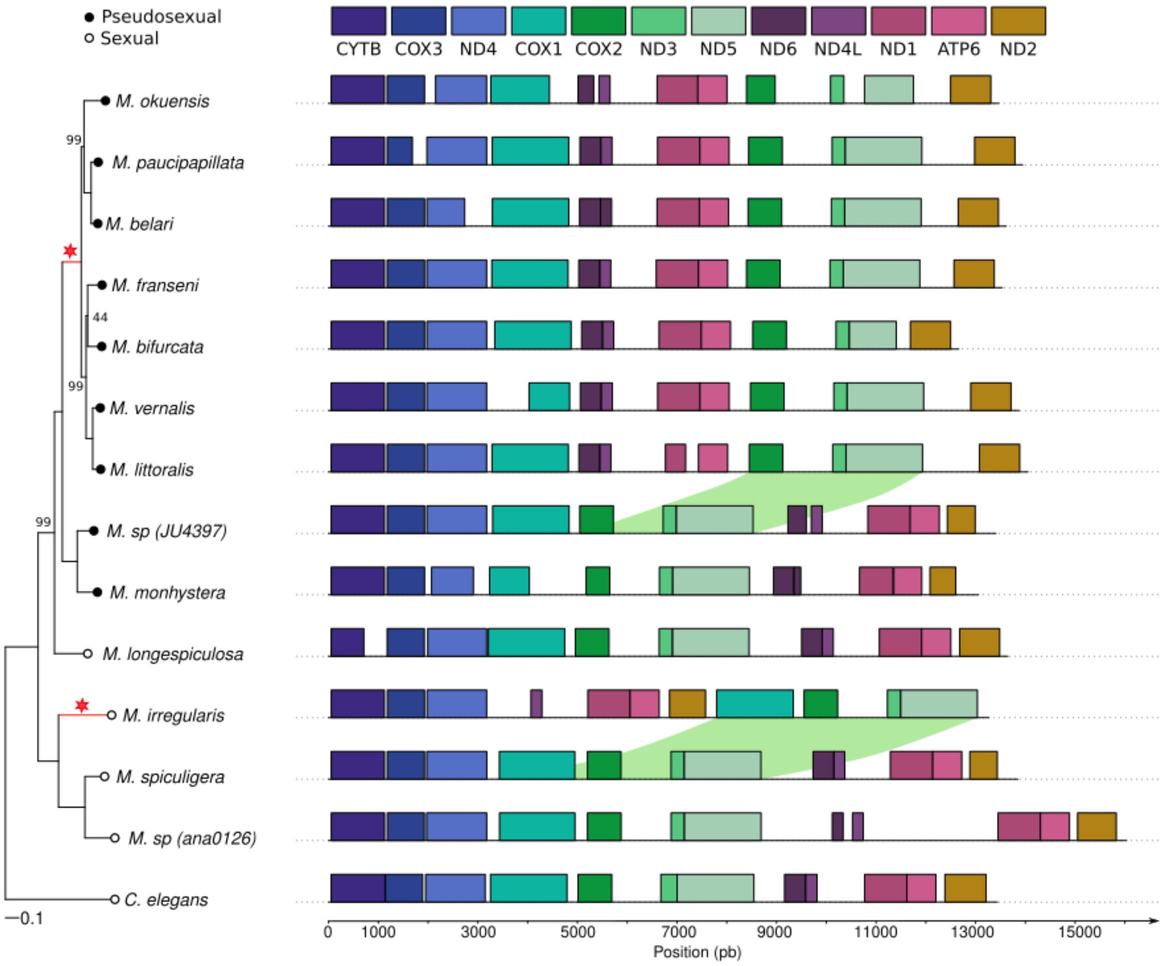


**Fig. S9. Two mitochondrial rearrangements have occurred in *Mesorhabditis*.** On the left, a phylogeny built from the mitochondrial genomes of our *Mesorhabditis* species, plus *C. elegans* as an outgroup, is shown. The two red stars label branches along which a mitochondrial rearrangement has occurred. Nodes are labelled based on mode of reproduction (filled and open circles). On the right, each mitochondrial gene is shown along the assembled mitochondrial genome of each species (coloured rectangles). The two rearrangements - which are translocations - are highlighted using green shadowing.

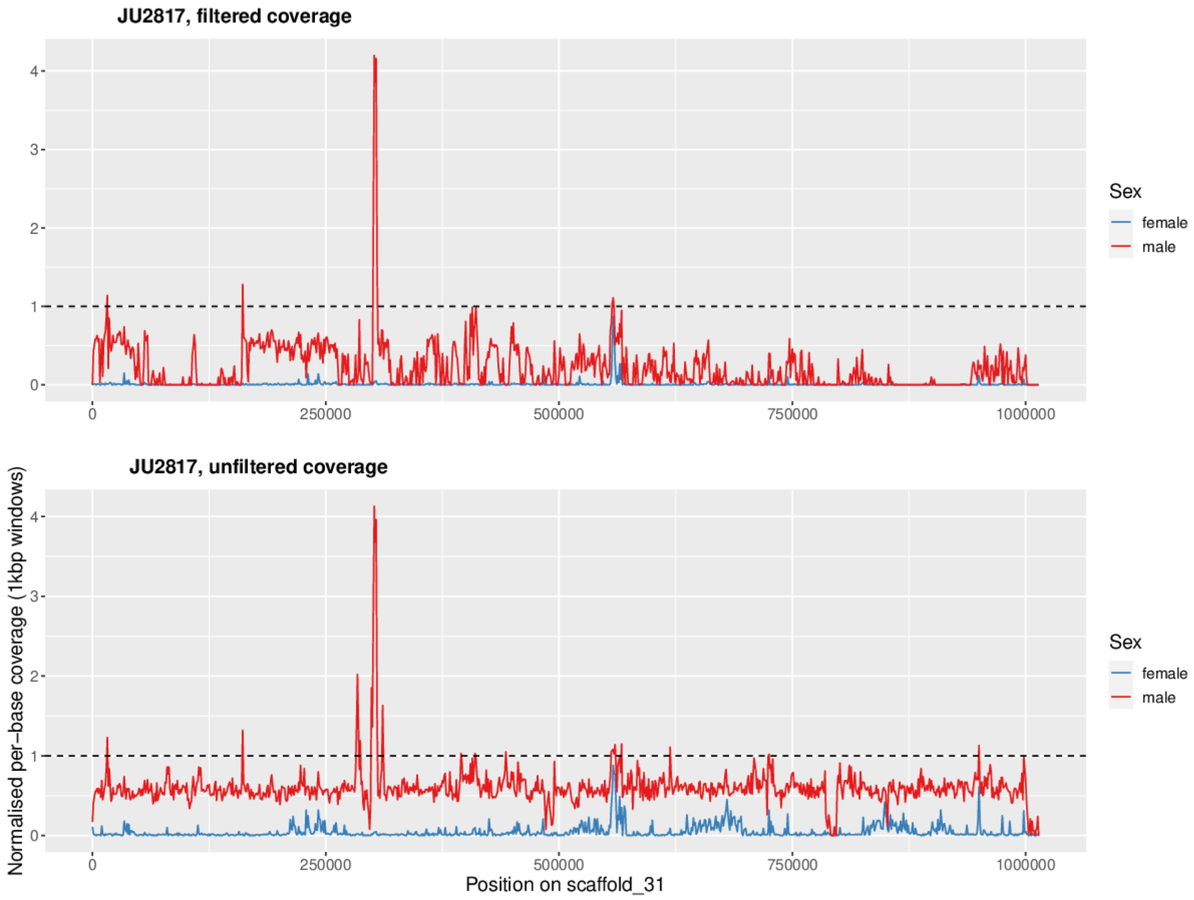


**Fig. S10**. **Filtered and unfiltered coverage on a Y-assigned scaffold in *M. belari*.** Normalised coverage for male and female DNA-Seq is shown (y-axis) along 1kbp non-overlapping windows of scaffold 31 (x-axis) of the *M. belari* assembled genome. The upper panel shows filtered coverage (MAPQ>=20 + reads are not marked as supplementary, secondary or duplicated), while the lower panel shows coverage but without the MAPQ filter (called 'unfiltered' here). Unfiltered coverage shows the expected haploid (values ~0.5) for males, but also shows some coverage in females. By contrast, filtered coverage shows almost no coverage in females, enabling more confidently assigning this scaffold to the Y. The peak in males probably corresponds to a duplicated region, while the region with filtered coverage in females (upper panel) may correspond to an assembly error.

| Scaffold 1 | Scaffold 2 | Contact type | Contact count | Ratio of inter- to intra-chromosomal contact | Contact counting approach |
|---|---|---|---|---|---|
| **scaffold_7** | **scaffold_16** | 3prime-3prime | 130 | 0.3 | pairix |
| scaffold_8 | scaffold_14 | 3prime-5prime | 100 | 0.265 | pairix |
| scaffold_14 | scaffold_23 | 5prime-5prime | 30 | 0.073 | pairix |
| scaffold_14 | **scaffold_16** | 5prime-5prime | 24 | 0.065 | pairix |
| scaffold_8 | **scaffold_16** | 3prime-5prime | 13 | 0.047 | pairix |
| scaffold_5 | **scaffold_16** | 5prime-5prime | 12 | 0.045 | pairix |

**Table S1: Top inter-scaffold Hi-C contact counts for *M. belari*.** Each row shows contacts computed between the extremities of two different scaffolds (first two columns), the kind of extremities the contacts are between (5prime or 3prime for start or end of the scaffolds - third column), the number of contacts (fourth column), the ratio of this number to the mean number of contacts inside each of the two extremities (fifth column), and the contact counting approach (see Methods for meaning). The top 10 inter-scaffold contacts, with a minimum of 10 contact counts, were selected and shown here. The X-assigned scaffolds are highlighted in bold. The highest inter-scaffold contacts are between the two X-assigned scaffolds of *M. belari*, however, when removing contacts occurring in duplicated regions of scaffolds, no contacts remain, as no rows with ‘contact counting approach’ set to ‘pairtools’ remain.

| Scaffold 1 | Scaffold 2 | Contact type | Contact count | Ratio of inter- to intra-chromosomal contact | Contact counting approach |
|---|---|---|---|---|---|
| scaffold_7 | scaffold_9 | 5prime-5prime | 54 | 0.137 | pairix |
| scaffold_9 | scaffold_10 | 3prime-3prime | 40 | 0.084 | pairix |
| scaffold_7 | scaffold_13 | 3prime-5prime | 34 | 0.081 | pairix |
| scaffold_9 | scaffold_13 | 5prime-3prime | 39 | 0.059 | pairix |
| **scaffold_8** | **scaffold_12** | 5prime-3prime | 127 | 0.028 | pairix |
| scaffold_3 | scaffold_13 | 3prime-3prime | 13 | 0.019 | pairix |
| scaffold_1 | scaffold_15 | 3prime-3prime | 36 | 0.001 | pairix |
| scaffold_1 | scaffold_4 | 3prime-5prime | 29 | 0 | pairix |
| scaffold_1 | scaffold_6 | 3prime-5prime | 23 | 0 | pairix |
| scaffold_1 | scaffold_3 | 3prime-5prime | 12 | 0 | pairix |
| scaffold_9 | scaffold_10 | 3prime-3prime | 14 | 0.121 | pairtools_mapq1 |
| scaffold_9 | scaffold_13 | 5prime-3prime | 12 | 0.063 | pairtools_mapq1 |
| **scaffold_8** | **scaffold_12** | 5prime-3prime | 48 | 0.034 | pairtools_mapq1 |
| **scaffold_8** | **scaffold_12** | 5prime-3prime | 45 | 0.032 | pairtools_mapq20 |

**Table S2: Top inter-scaffold Hi-C contact counts for *M. monhystera*.** For the meaning of the columns, see table above. The X-assigned scaffolds are highlighted in bold. Contacts between the two X-assigned scaffolds are consistently high. In particular, when filtering out contacts occurring in duplicated regions of scaffolds (i.e. looking only at rows with ‘contact counting approach’ set to ‘pairtools’), and with this filtering threshold set to a high value (MAPQ of 20, meaning the probability that a contact is a false positive is <~1%), only contacts between the two X-assigned scaffolds remain.

| | X | Y | autosome |
|---|---|---|---|
| #orthogroups | 1521 | 56 | 10927 |
| # uniquely assigned orthogroups (% of all orthogroups) | 1415 (93%) | 12 (21%) | 10223 (94%) |
| #genes | 1549 | 139 | 13857 |

| | | | |
|---|---|---|---|
| #genes/orthogroup | 1.02 | 2.48 | 1.27 |
| gene density per kbp | 0.102 | 0.018 | 0.090 |

**Table S3: Gene- and gene family-level statistics for *M. belari*.** Columns show X-, Y- and autosome-assigned genes. An orthogroup is a family of genes as defined using OrthoFinder (see Methods), and can include genes assigned to both X, Y and autosomes. #orthogroups is a count of orthogroups with at least one X, Y or autosome-assigned gene member. # uniquely assigned orthogroups counts orthogroups with only members assigned to X, Y or autosomes.

| | X | Y | autosome |
|---|---|---|---|
| #orthogroups | 1677 | 79 | 10189 |
| # uniquely assigned orthogroups (% of all orthogroups) | 1567 (93%) | 19 (24%) | 9771 (96%) |
| #genes | 1721 | 246 | 13488 |
| #genes/orthogroup | 1.03 | 3.11 | 1.32 |
| gene density per kbp | 0.101 | 0.014 | 0.090 |

**Table S4: Gene- and gene family-level statistics for *M. monhystera*.** For table legend, see table above.

| | *M. belari* | *M. monhystera* |
|---|---|---|
| X: span, %covered by repeats | 15.1 Mbp, 22.2% | 17.1Mbp, 28.0% |
| Y: span, %covered by repeats | 7.8Mbp, 70.2% | 17.0Mbp, 88.9% |
| autosomes: span, %covered by repeats | 153.2Mbp, 34.0% | 149.4Mbp, 32.7% |

**Table S5: The Y chromosome is enriched for repeats compared to the X and autosomes in the pseudosexual species**. For the two pseudosexual species, the span (total length) of all X, Y and autosomal-assigned scaffolds is shown (rows), together with the fraction of those scaffolds covered by repeated elements (Transposable elements, Satellites and other uncharacterised repeated elements) as predicted using EarlGrey (see Methods).
